# Identification of cellular and molecular risk signatures for progression to late-stage age-related macular degeneration using the 9-step Minnesota Grading System

**DOI:** 10.1101/2025.07.10.664209

**Authors:** Chia-Ling Huang, Yubin Qiu, John T Demirs, Xiaoqiu Wu, Michael Twarog, Omar Delgado, Maura Crowley, Natasha Buchanan, Nhi Vo, Lin Fan, Yanqun Wang, Junzheng Yang, Thomas R. Vollmer, Garrett Klokman, Sandra Jose, Jorge Aranda, Ganesh Prasanna, Stephen Poor, Cynthia L. Grosskreutz, Timothy W. Olsen, Christopher W. Wilson, Magali Saint-Geniez, Sha-Mei Liao

## Abstract

Age-related macular degeneration (AMD) is a complex multifactorial disease, and the molecular mechanisms underpinning the progression of intermediate AMD to geographic atrophy are not fully understood. To better understand mechanisms driving progression, we performed bulk RNA sequencing on dissected macular and peripheral RPE/choroid and neural retina tissue from postmortem human eyes graded using the 9-step Minnesota Grading System (MGS). Binning of intermediate AMD cases into three distinct groups (AMD3L, AMD3M, AMD3H) based on the 5-year risk of progression enabled identification of distinct gene and pathway changes associated with progression to late-stage disease. Identified changes in gene expression were validated using ELISA or histological methods. RPE-specific genes and lipid metabolic pathways showed a transient increase in AMD3L followed by a pronounced decrease in AMD3H. In AMD3H, immune response genes such as *C3*, *TREM2*, and *OLR1* were upregulated when compared to AMD3L samples, as well as genes specific to Müller glia/astrocytes *(NGFR, SPP1, GPX3*). Our findings support complement inhibition as a promising therapeutic option for slowing conversion to advanced AMD and identify macrophage and Müller/astrocyte genes as potential cell types to target in AMD. Further, we demonstrate the value of combining emerging, outcomes-based, clinically relevant grading systems with profiling technologies to generate new insights into ocular diseases.

**Highlights:**

- Intermediate AMD (AMD3) can be further divided into 3 stages - AMD3L (low risk), AMD3M (intermediate risk), AMD3H (high risk) – using the MGS9 grading system based on the risk of disease progression to late AMD.
- RNA sequencing of the macular RPE/choroid shows opposing changes in gene expression of multiple biological pathways for both RPE and immune cells between AMD3L and AMD3H stages.
- In the macular neural retina, most biological pathways were downregulated in AMD2 (early AMD) but upregulated in AMD4 (late AMD) compared to AMD1 (non-AMD control).
- Molecular and cellular signatures associated with a high risk of progression to AMD4 include activation of complement C3, two subtypes of macrophages expressing either TREM2 or OLR1, and Müller glia/astrocytes as evidenced by the upregulation of GFAP and NGFR.
- Understanding the roles of these high-risk associated genes in AMD progression will facilitate the development of new treatments that prevent or delay the irreversible central vision loss in AMD patients.

## 1. Introduction

Age-related macular degeneration (AMD) is a leading cause of blindness, with the projected number of people suffering from AMD expected to rise from 196 million in 2020 to 288 million in 2040 ^1^. AMD is a chronic, slow progressing, and degenerative eye condition that gradually reduces central vision because of macula thinning caused by irreversible loss of retinal pigment epithelium (RPE) and photoreceptors. There are two forms of the advanced stage of the disease: geographic atrophy (GA), the end stage of dry AMD, and wet/neovascular AMD (nAMD). Although anti-VEGF therapies are effective for nAMD patients, 96% of them will eventually progress to GA, resulting in irreversible loss of central vision in a 7-year follow up period ^2^. The detailed pathophysiology of dry AMD remains unclear due to the multifactorial nature of the disease as described in human genome-wide studies ^3–7^.

Developing treatments for GA has proven challenging, with only modest efficacy seen with complement-targeting therapies so far ^8^. This raises the possibility that treating AMD at an earlier stage of the disease might be more effective in halting progression to GA or nAMD. However, there are currently no approved treatments available for intermediate AMD patients. An oral complement factor B inhibitor, Iptacopan, is in phase 2 clinical trials with the aim to prevent conversion of early or intermediate AMD eyes to new incomplete RPE and outer retinal atrophy (iRORA) ^9^ or late AMD (NCT05230537). Profiling the molecular changes that occur prior to GA/nAMD would provide valuable information about the specific cell types and biological pathways involved in disease progression. Identifying signatures associated with progression could facilitate the discovery of new therapeutic targets, ultimately helping to develop more effective interventions that prevent or delay the irreversible central vision loss by preserving the health of RPE and photoreceptors.

Significant progress has been made in understanding the pathogenic mechanisms of AMD. Technologic areas of advancement include imaging, genetics, epigenetics, transcriptomics, proteomics, and metabolomics ^10–21^. Most gene expression studies conducted so far have utilized a classification system consisting of 3 to 4 stages of the disease: AMD1 (normal/non-AMD), AMD2 (early AMD), AMD3 (intermediate AMD), and AMD4 (GA and/or nAMD) ^22–24^. However, this approach has posed challenges in capturing high-risk signals for progression to late-stage AMD, as the 5-year risk of progression to GA/nAMD in AMD3 is highly heterogenous and can range from 2 to 53% ^25^. In this study, we employed the 9-step Minnesota Grading System (MGS9) to sub-stratify post-mortem AMD3 eyes into three groups based on their low (AMD3L), intermediate (AMD3M), or high risk (AMD3H) of progression to the end stage of AMD. We conducted transcriptional profiling of RPE/choroid and neural retina tissues at every stage of AMD to identify stage-associated changes in gene expression, and validated genes associated with a higher risk of progression to GA using additional assays. The genes identified in this study may serve as cellular and molecular signatures for the risk of AMD progression to GA and suggest potential therapeutic targets for intermediate AMD.

## 2. Methods

### 2.1 Post-Mortem Donor Eye Collection and AMD Severity Grading

Post-mortem human eyes were procured by the Lions World Vision Institute, (LWVI, formerly Lions Eye Institute for Transplant & Research/LEITR; Tampa FL) with consent of donors or donors’ next of kin and in accordance with the Eye Bank Association of America (EBAA) medical standards, US/Florida law for human tissue donation, the Declaration of Helsinki and FDA regulations, the guidelines of the ARVO Best Practices for Using Human Eye Tissue in Research (Nov 2021), and Novartis guidelines regarding research using human tissues. Detailed methods and reagents for post-mortem eye collection were described previously ^26^. Briefly, all study samples were collected and preserved within 6 and a half hours or less post-mortem. The anterior chambers of collected donor eyes were removed by making a circular incision parallel to and about 4 mm below the limbus. The remaining posterior eye tissues were dissected into four distinct regions and flash frozen for RNA/protein analysis: the macular RPE/choroid, peripheral RPE/choroid, macular neural retina, and peripheral neural retina. Macular tissue was isolated from a 6-mm diameter macular punch and the entire peripheral tissue was collected. High-resolution, stereoscopic color fundus photographs of the posterior eye were taken using a dissection microscope and camera both before and after removal of the neural retina. AMD disease stages were assigned after reviewing fundus images using criteria from the Minnesota Grading System (MGS) for assignment into 4 stages (MGS-4; normal/unaffected controls = AMD1, early = AMD2, intermediate = AMD3, and end-stage AMD = AMD4) and separately divided into 9 stages (MGS-9) as described ^27,28^.

### 2.2 Bulk RNA-seq

Total RNA was isolated from homogenized RPE/choroid and retina tissues using a RNeasy Plus Mini kit (Qiagen). RNA integrity (RIN) was assessed using the 18S/28S rRNA bands in capillary electrophoresis (Bioanalyzer, Agilent Technologies, Santa Clara, CA). For the bulk RNAseq study, the peripheral and macular RPE/choroid libraries were generated manually with an Illumina Stranded mRNA Prep Ligation kit (cat# 20040534; user guide # 1000000124518 v01), using 100 ng and 60 ng total RNA input for each sample, respectively. One µl of 1:200 ERCC RNA Mix 1 was added to each sample (Ambion, 4456740). Universal Human Reference RNA (Agilent, 740000), 100 ng and 60ng, respectively, was used as a positive control and water with/without ERCC were used as a negative control. Libraries were checked for quality using an Agilent TapeStation D1000 and sequenced on an Illumina NovaSeq 6000 platform with 51 bp paired-end reads. The peripheral and macular retina libraries were generated with an Illumina TruSeq® Stranded mRNA Sample Preparation kit (part # 15031057, Rev. E) on Hamilton STAR, using 200 ng total RNA input for each sample. Two µl of 1:200 ERCC RNA Mix 1 was added to each sample (Ambion, 4456740). Two hundred ng of Universal Human Reference RNA (Agilent, 740000) was used as positive control and water was used as negative control. All other quality control and sequencing steps were performed identically as to the RPE/choroid libraries. Read alignment and quantification were conducted using STAR (v2.6.0), RSEM (v1.2.22) and the human reference genome downloaded from GENCODE (GRCh38.p10). Samples with sequence read counts less than 10M were excluded from the analysis due to low sequencing depth. Principal component analysis was performed to assess intra-group and inter-group sample variations. Differential expression analysis was conductedusing edgeR-limma-voom workflow ^29^ and adjusted for sex. All data processing and analyses were performed using custom python and R scripts. The raw fastq files have been deposited in Gene Expression Omnibus (GEO)^30^ (accession number pending). Functional gene set enrichment analysis (GSEA) was carried out using a differential gene list ranked by t-score for each comparison ^31,32^. Hallmark and REACTOME gene sets from MSigDB v7.3 were utilized first for pathway enrichment analysis. The RPE and macrophage marker genes were from MSigDB C8: cell type signature gene set HU_FETAL_RETINA_RPE and ZHENG_CORD_BLOOD_C9_GRANULOCYTE_MACROPHAGE_PROGENITOR ^33,34^.

### 2.3 Single-nucleus RNA-seq (snRNA-seq)

A reference single-nuclear expression library was generated from post-mortem human tissue, from either (1) a 6 mm macular RPE-choroid biopsy punch or (2) posterior eye sections from whole globes [Orozco et al]. For whole globes, a donor eye was enucleated and flash frozen in liquid nitrogen 4 hours and 34 minutes post-mortem. The anterior segment of the globe and vitreous were removed. The posterior eye cup was embedded in OCT (Tissue-Tek), flash frozen, and cryo-sectioned from sclera to retina. Three 60 µm sections were collected for snRNA-seq profiling of the major cell types in the back of eye. Nuclei were extracted from RPE-choroid biopsy punch or cryosections using a modified version of the ‘Frankenstein’ protocol described by Martelotto et al. ^35^, stained with DAPI (Thermo Fisher Scientific), and sorted by fluorescence activated cell-sorting (FACS). Debris were removed based on forward and side-scatter, and cells were selected for events with 2n DNA content as previously described ^23^. Purified nuclei were counted using Acridine orange and propidium iodide dye (Nexcelom) with an epifluorescent microscope and prepared for single nucleus RNA-seq using a 3’ gene expression kit (v3) from 10X Genomics, following manufacturer’s recommended protocols. Library quality was measured via a High Sensitivity DNA kit using a Bioanalyzer (Agilent). Sequencing libraries were pooled and sequenced using an Illumina NovaSeq 6000 per manufacturer’s instructions.

Read alignment and generation of feature barcoded matrices was performed using CellRanger v3.02 and the human GRCh38 reference genome. Single nuclei RNA-seq quality control, normalization and clustering were run using the Seurat v3 R package ^36^ under R environment 3.6. Cells with nFeatures ≥ 7447 (99^th^ percentile) or ≤ 439 (1^st^ percentile) were excluded from the analysis to avoid doublets or empty droplets. Low quality cells with a percentage of mitochondrial transcript reads ≥ 20% and a percentage of ribosomal reads ≥ 2% were excluded to avoid apoptotic or lysing cells^37^. After quality control, 15,613 cells were used for follow-up analysis. After clustering, clusters were annotated based on the expression of canonical markers (**Supplementary Table 6**). The count matrix was deposited in GEO (accession number pending).

### 2.4 Deconvolution of bulk RNA-seq with cell-type specific signatures

Cell type deconvolution of bulk RNA-seq data was performed using snRNAseq data as a reference. The snRNAseq data was downsampled to 200 cells per cell type to improve computational time. Analysis was performed using the MuSiC v0.2.0 algorithm with default parameters ^38^. Deconvolution analysis was performed for each individual bulk sample.

### 2.5 RNAscope/in situ hybridization and immunohistochemistry

Donor eyes for histology were processed, sectioned, and stained with hematoxylin and eosin (H&E) as previously described ^26^. AMD grading of H&E-stained histological sections was determined following published criteria and as described in the text ^27,39,40^.

Detection of mRNA by RNAscope was performed using Advanced Cell Diagnostics probes (Advanced Cell Diagnostics/ACD, Newark, CA) against human *C3* (#430709), human *OLR1* (#517529), human *GPX3* (#470599), human *SPP1* (#420109), human *NGFR* (#406399), and human *TREM2* (#420499). Probes were hybridized and detected using the RNAscope® VS Universal AP Reagent Kit (ACD), comprised of the RNAscope VS Universal AP Detection Reagents (PN 323260), the RNAscope VS Accessory Kit (PN 320630), and the RNAscope Universal VS Sample Preparation Reagents v2 (PN 323740) for RNA in situ hybridization (ISH) and the single-plex chromogenic RED assay designed for use with Discovery Ultra automated IHC/ISH slide staining systems by Ventana Medical Systems. Slides were scanned using an Aperio X1 brightfield scanner (Leica Biosystems 21440 W, Deer Park, IL) and transferred to HALO image analysis software (Indica Labs, Albuquerque, NM) for further analysis.

For immunohistochemistry, detection of IBA1 (Fujifilm WAKO 019-19741, rabbit polyclonal, dilution 1:1000) and glial fibrillary acidic protein (GFAP, Proteintech; 16825-1-AP, rabbit polyclonal, dilution 1:2000) was done on FFPE sections using a Ventana Ultra with standard alkaline phosphatase (AP) chromogenic detection (Roche, ChromoMap Blue 760-161). Co-detection of human *C3* mRNA with IBA1 was conducted using a LeicaBond RX system. Probes for human *C3* (#430708) and the RNAscope-RED detection kit (#322150) were from ACD. *C3* mRNA staining was performed first, followed by IBA1 protein detection with green chromogen (#DC9913, Leica Biosystems). Slides were scanned and analyzed as described above.

### 2.6 TREM2 MSD/ELISA

Human donor tissues were dissected, weighed, and processed as previously described ^26^. Total TREM2 protein levels were measured using a TREM2 Duoset (R&D Systems, Minneapolis, MN) ELISA kit for MSD-based assays (MSD, Rockville, MD) and read on a Sector S 600 imager. TREM2 concentrations in fluids (ng/ml) or tissue homogenates (ng/g, normalized to weight of dissected tissue) were converted to molar concentrations based on the predicted molecular weight of TREM2 (25.4 kDa).

### 2.7 Statistics

Prism software (GraphPad, www.graphpad.com) was used for statistical analysis. Data are presented as mean ± standard error of mean (SEM). Comparisons of means for levels between groups were generated using Student’s t-test (unpaired, 2-tailed)

## 3. Results and Discussion

### 3.1 Stratification of intermediate AMD/AMD3 into three groups based on the risk of progression towards late or advanced AMD using the 9-step Minnesota grading system

The Minnesota Grading System initially defined AMD in a four-stage disease scale (MGS4), primarily based on drusen size assessed from post-mortem color fundus images of the macular area ^27^. This scale includes AMD1 (normal/non-AMD), AMD2 (early AMD), AMD3 (intermediate AMD), and AMD4 (A and/or neovascular AMD). Studying molecular changes that correlate with AMD disease progression has been hampered by heterogeneity in AMD3 samples, as the 5-year risk of disease progression from AMD3 to AMD4 is highly variable and ranges from 2% to 53% ^25^. A refined Minnesota Grading System (MGS9) was developed based on total drusen area and hypo/hyper-pigmentation changes in the macula, which further divides the AMD3 group into 7 steps (MGS9-3 to MGS9-9) ^28^. The MGS9 system is designed to parallel the clinically utilized Age-Related Eye Disease Studies (AREDS) system, providing a meaningful correlation to the known risk of AMD progression. We collected 102 post-mortem human eyes that were initially graded using the MGS4 system. Fifty-six of these eyes were graded as AMD3 using MGS4; these were subsequently re-graded using MGS9 to obtain better sample stratification. We then grouped the seven stages of AMD3 in the MGS9 system into three bins. We defined MGS9-3 to MGS9-5 as AMD3L, which has a low risk (2-6%) of progressing to advanced disease progression in five years. According to fundus features in postmortem eyes, AMD3L maculas typically have a smaller drusen area and no pigmentary changes. The medium risk group, AMD3M, encompassed stages MGS9-6 to MGS9-7, with a 14-28% risk of progression. AMD3M is characterized by intermediate ranges of drusen area and pigmentary changes. AMD3H spans stages MGS9-8 to MGS9-9 and has the highest 5-year risk of progression (47-53%). This grouping is associated with the largest drusen area, hypo/hyper-pigmentation changes, and/or non-central geographic atrophy. Mapping of the MGS4, MGS9, and our groupings to the 5 year-risk of progression is shown in **Figure 1A**. Examples of fundus images taken in the dissected macular area after removal of the retina for AMD1, AMD2, AMD3L/M/H, and AMD4 can be seen in **Figure 1B**, with accompanying donor information (**Supplemental Table 1**). In this figure, 4-step MGS grades are annotated as MGS4, and 9-step MGS grades are described as MGS9, followed by the relevant numbers on each scale.

**Figure 1.**
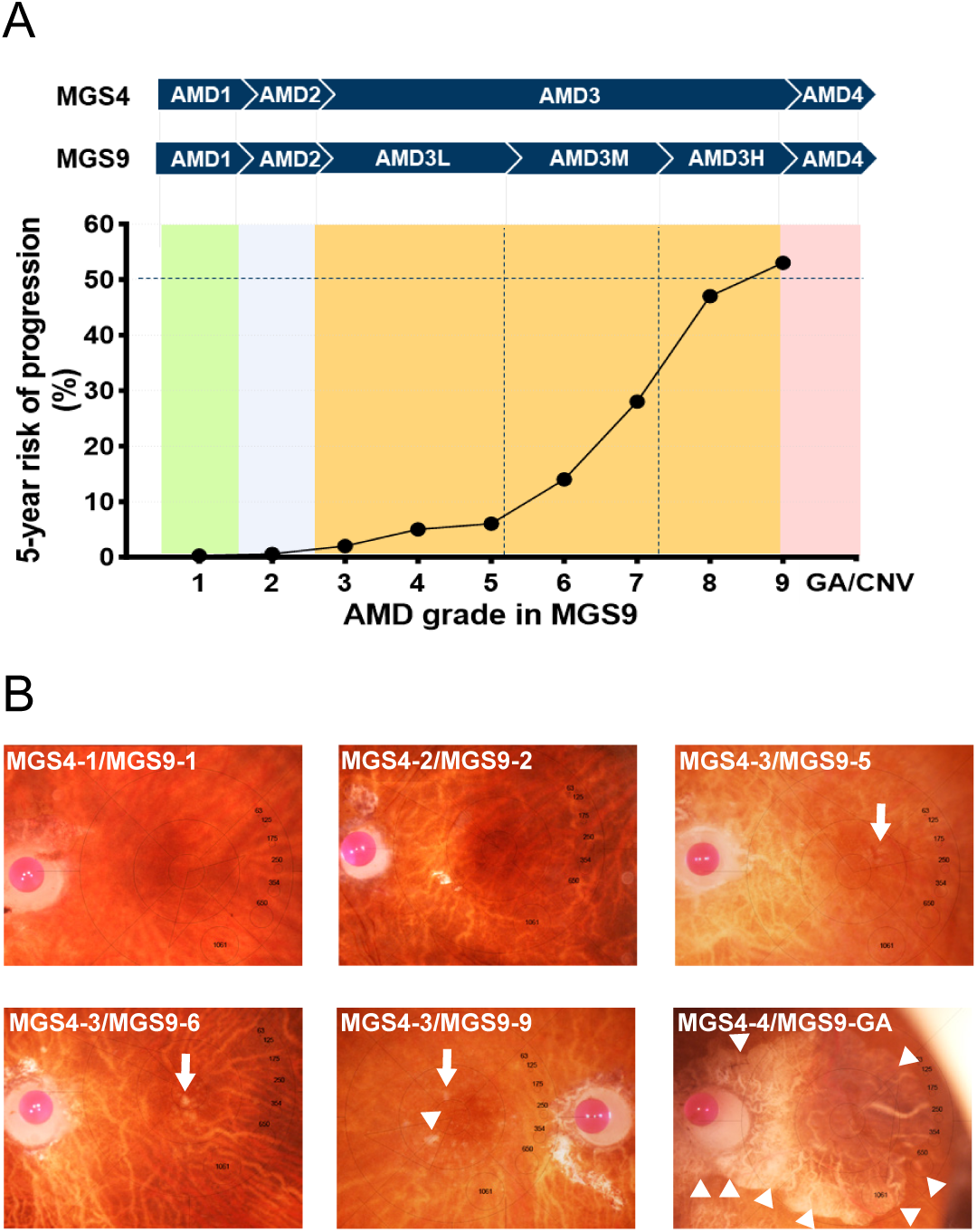
Stratification of intermediate AMD into 3 groups based on the risk of progression to late/advanced AMD using the 9-step MGS. **A**. Correlation of MGS9 AMD disease severity grading of postmortem eyes with the risk of progression to advanced AMD ^25,28^. MGS4-1 is equivalent to MGS9-1, MGS4-2 is similar to MGS9-2, MGS4-3 spans the MGS9-3 to −9 grades, and, MGS4-4 represents geographic atrophy (GA) or/and neovascular AMD (nAMD). Based on the 5-year risk of progression to late AMD, the MGS4-3 can be further divided into 3 groups: AMD3L, which includes MGS9-3 to −5; AMD3M, which includes MGS9-7 and −8; and AMD3H, which includes MGS9-8 and −9. **B**. Representative color fundus images of the macular RPE/choroidal area from post-mortem donor eyes, taken after the retina was removed. The macula area is denoted by a 6 mm circle, while the smallest 1 mm circle signifies the fovea. Various drusen sizes are marked by circles numbered from 63 to 1061 µm, serving as reference points. Arrows indicate typical drusen, and arrowheads indicate RPE atrophy/loss. Images include AMD1, a normal macula; AMD2, showing minimal drusen; AMD3L, an MGS5 eye, with several drusen; AMD3M, an MGS6, showing moderate drusen area and minimal pigment changes; AMD3H, an MGS9, with more abundant drusen and non-foveal RPE atrophy; the AMD4 eye features geographic atrophy characterized by a significant large area of RPE loss, outlined by arrowheads. All eyes depicted are from left eyes (OS), except the AMD3H, which is a right eye (OD).

### 3.2 MGS9-based stratification identifies transcriptome changes in metabolic and immune pathways in the macular RPE/choroid

Previously published bulk RNA-seq studies were conducted with postmortem eyes classified using four-step AMD grading ^22–24^. We asked if further substratification of bulk RNA-sequenced samples could unveil additional gene and cell signatures indicative of disease progression from early/intermediate to late/advanced stages of AMD. RNA was prepared from dissected macular and peripheral tissues of the RPE/choroid and neural retina (refer to **Supplemental Table 2A** for the list of samples classified by MGS4, **Supplemental Table 2B** for the list of samples classified by MGS9, **Supplemental Table 3** for the list of individual donor information). After bulk RNA sequencing and data processing, we first evaluated differential gene and pathway expression by region and dissected tissue. When all groups were compared against AMD1 as a baseline, statistically significant changes at the gene level were only observed in the AMD4 group (**Supplemental Table 4**). Higher numbers of differentially expressed genes (DEGs) were detected in the macular tissues compared to the peripheral tissues (**Supplemental Table 4**), consistent with AMD being a macular disease. Since we did not detect DEGs in intermediate AMD/AMD3, perhaps owing to its subclinical phenotype and changes potentially limited to a small proportion of cells, we prioritized gene set enrichment analysis (GSEA) to initially assess changes in biological processes ^41^.

GSEA pathway analysis of our macular RPE/choroid RNAseq data, comparing all groups to AMD1 using Hallmark genesets, was able to identify statistically significant pathway changes and categorize them into several clusters over AMD progression, using both the MGS4 (**Figure 2A**) and MGS9 (**Figure 2B**) classifications. Notably, more pronounced changes were observed when subclustering AMD3 into AMDL, AMDM, and AMDH groups. Using this system, we identified four clusters of gene expression (**Figure 2B**). Cluster 1 (purple) was comprised of pathways that increase in all AMD stages compared to AMD1; notable pathways include metabolic signaling and some interferon-related pathways. Cluster 2 (green) consisted of pathways (bile acid metabolism, Notch, KRAS) that were upregulated in early AMD2/AMD3L and then reduced in AMD3H. Cluster 3 (blue) showed opposite patterns, with key pathways such as hypoxia, TNFα, and IL6 signaling showing downregulation in AMD2 and AMD3L, followed by upregulation in AMD3H. Lastly, pathways in Cluster 4 (red) showed consistent downregulation when compared to AMD1. The complex patterns emphasize distinct or opposing pathway changes in temporal progression from AMD3L to AMD3H, which were not as readily observed when combining all samples into an intermediate AMD3 group.

**Figure 2.**
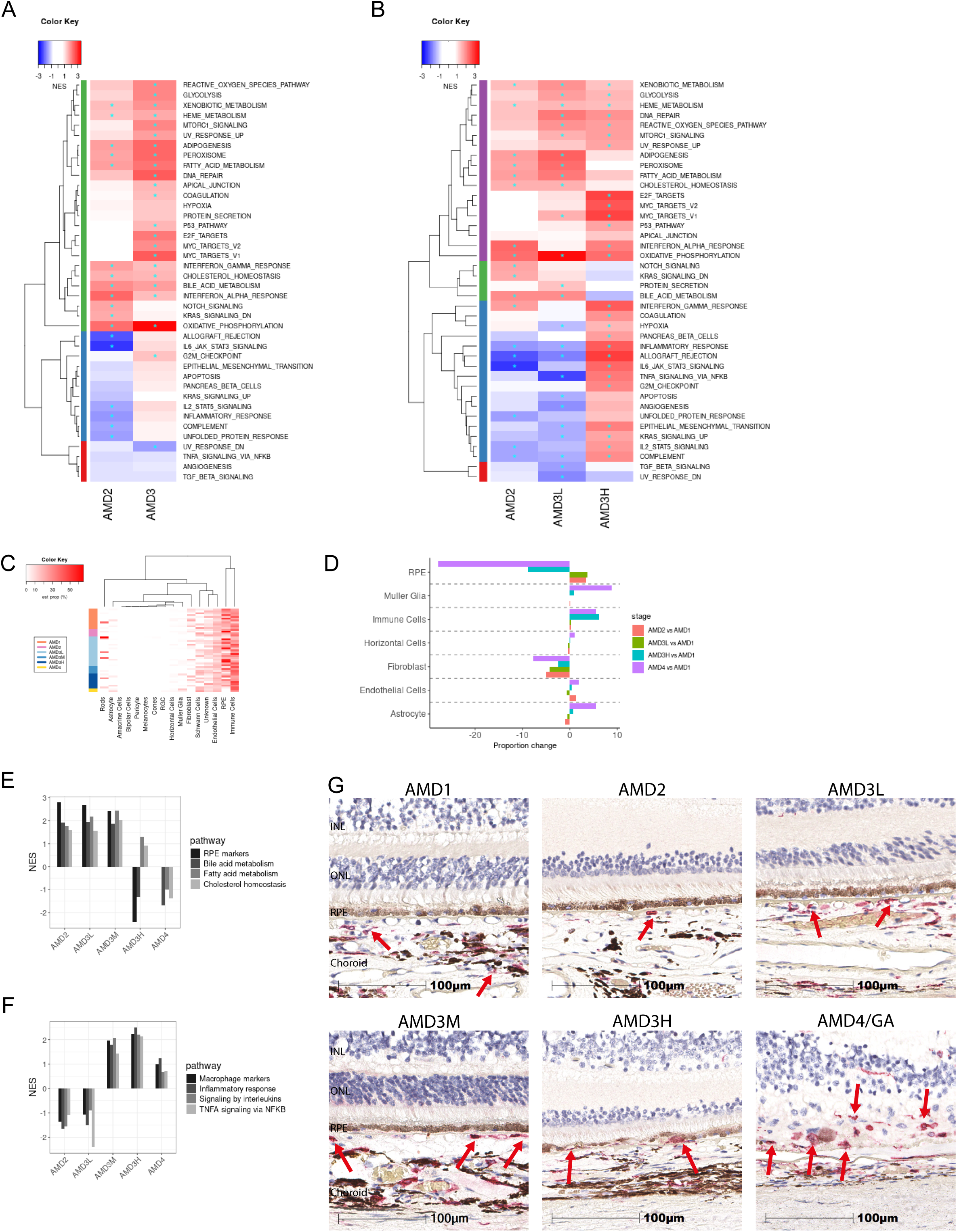
Stratification of bulk RNAseq data from AMD3 sample into low- and high-progression risk groups identifies progression-associated pathways and changes in cell proportions. **A**. Clustered heatmaps of normalized enrichment scores (NES) analyzing GSEA Hallmark genesets across RNAseq data from the macular RPE/choroid, comparing AMD2 and AMD3 to AMD1 in the left panel, and AMD2, AMD3L, and AMD3H to AMD1 in the right panel (**B**). Upregulated pathways (positive NES value) are displayed in red and downregulated ones (negative NES value) in blue. * represents a false discovery rate (FDR) lower than 0.05. The hierarchical cluster groups in the dendrogram, indicated by colored bars on the y-axis, represent pathways with similar changes in AMD progression. Sample numbers: AMD1 = 11; AMD2 = 4, AMD3 = 28; AMD3L = 16, AMD3H = 8. **C.** Heatmap of cell proportions in macular RPE/Choroid across AMD1 to AMD4. Cell proportions in the bulk RNAseq data were inferred using the MuSiC algorithm ^38^, utilizing cell-type specific gene signatures from a single nucleus RNA-seq dataset (**Supplemental Figure S2**). The term “est prop (%)” refers to the estimated proportion of a given cell type. Among all cell types, immune cells, RPE, and endothelial cells represent the highest proportions. **D**. Bar graph of cell-type proportion changes across AMD2, AMD3L, AMD3H, and AMD4 as compared to AMD1 in selected cell-types. RPE cell proportion increased in AMD2 and AMD3L but was reduced in AMD3H and AMD4. Immune cell proportion was increased in AMD3H and AMD4. Neural retinal cell types such as Müller glia, astrocytes, and horizontal cells were only detected in AMD4 RPE/choroid samples. **E.** NES of RPE cell-specific gene signatures shows a general concordance with NES of lipid metabolic pathways (bile acid, fatty acid, cholesterol metabolism) across AMD stages when compared to AMD1. **F.** NES of macrophage-specific gene markers show concordance with immune response signaling pathways across AMD stages when compared to AMD1. **G.** Representative images of IBA1 staining of outer retina/choroid sections across AMD grades. In AMD1, most IBA1^+^ cells (red arrows) are located in the deep choroid and are not proximal to the choriocapillaris/BrM/RPE. In AMD2, some isolated IBA1^+^ cells can be observed within the CC space. In AMD3L and AMD3M, more IBA1^+^ cells are located in the CC space. In AMD3H, IBA1^+^ cells are detected in the sub-RPE space. In AMD4, numerous IBA1^+^ cells are observed around the GA lesion area and in the neural retina where most RPE cells and PR have degenerated.

Cell composition analysis of bulk RNA-seq data with the MuSiC v0.2.0 ^38^, using a single nucleus RNA-seq of sectioned posterior eye as cell-type specific reference (**Supplemental Figure S1, Supplemental Table 5**), indicates that the major RNA components in macular RPE-choroid bulk RNA-seq originate from endothelial cells, RPE, and immune cells (**Figure 2C**). Based on mRNA assessments, the RPE cell proportion increased in AMD2-3L, decreased in AMD3H, and further decreased in AMD4 (**Figure 2D**). This is largely consistent with RPE loss/degeneration defining AMD grading, supported by histological evidence of RPE atrophy as well as changes in RPE gene signature and lipid pathways (**Figure 2E**). Multiple lipid metabolic pathways in RPE cells are required for the constant clearance of the lipid-rich photoreceptor outer segments. The immune cell proportion was increased in AMD3H and AMD4 but was lower in earlier stages. This correlates with the peak levels of macrophage gene signature and inflammatory/cytokine pathways in AMD3H (**Figure 2F**) suggesting that the inflammatory-immune response may play a prominent role in the risk of disease progression to late AMD. This is supported by histological observations of choroidal IBA1+ macrophages positioned closely to the RPE layer in AMD2 and AMD3L and proximal to the sub-RPE space in AMD3H (**Figure 2G**). In contrast, very few IBA1+ cells were detected in the macular photoreceptor/RPE/choriocapillaris of AMD1 donors, suggesting that activation of choroidal macrophages or recruitment of circulating monocytes occurs in the earlier stages of AMD. Additionally, mRNAs specific to astrocyte/Müller and horizontal cells were also detected in AMD4 RPE/choroid (**Figure 2C-D**), likely due to PR/RPE degeneration leading to the adhesion of glial and retinal cells to RPE cells or Bruch’s membrane (BrM).

### 3.3 In the macular neural retina, most pathways were downregulated in AMD2 but upregulated in AMD4

GSEA pathway analysis of macular neural retina RNAseq data with Hallmark gene sets showed that most pathways were downregulated in AMD2 compared to AMD1, when using the MGS4 grading system, with several pathways showing upregulation in AMD3 (**Figure 3A**). When substratification of AMD3 samples into L/M/H was conducted, similar clusters emerged as in the macular RPE/choroid, when comparing pathway expression to AMD1 (Figure 3B?). Cluster 1 (purple) showed downregulation of pathways in AMD2 and upregulation in AMD3L and AMD3H, including angiogenesis and TGFβ signaling. Cluster 2 (green) was comprised of two pathways (interferon alpha and E2F targets) that showed trends or statistically significant upregulation in AMD2, AMD3L and AMD3H. Cluster 3 (blue) was comprised of pathways that oscillated between downregulated in AMD2, up in AMD3L, and downregulated again in AMD. Cluster 4 (red) consisted of pathways that were downregulated in AMD2 and AMD3L, and were upregulated in AMD3H. Cell composition analysis suggests that RNA from rod photoreceptors accounted for approximately 80% of the total RNA in AMD1, AMD2 and AMD3 samples, consistent with rods being the predominant cell type in the macular neural retina (**Figure 3B**). The relative proportion of rod photoreceptors progressively decreased from AMD3L to AMD4 (**Figure 3C**), consistent with rod photoreceptor loss in macular degeneration. In contrast, the proportion of Müller glia/immune cells was highest in AMD4. Müller/astrocyte activation in AMD4 was confirmed by GFAP staining (**Figure 3D**). GFAP, which is primarily expressed in astrocytes, was present in the ganglion cell layer in AMD1 and in the non-atrophic area of an AMD4 tissue section (**Figure 3E**). In the AMD transition zone from the non-atrophic to the atrophic area, GFAP processes gradually expand through the entire neural retina, eventually reaching the RPE and BrM in the atrophic area of AMD4 where there is severe RPE/PR loss (**Figure 3E**). As expected, IBA1+ immune cells, likely microglia, were detected in the inner macular retina in AMD1 but appear to migrate and accumulate to the PR layer and subretinal space in AMD3H/AMD4 (**Figure 2G, Supplemental Figure 3**).

**Figure 3.**
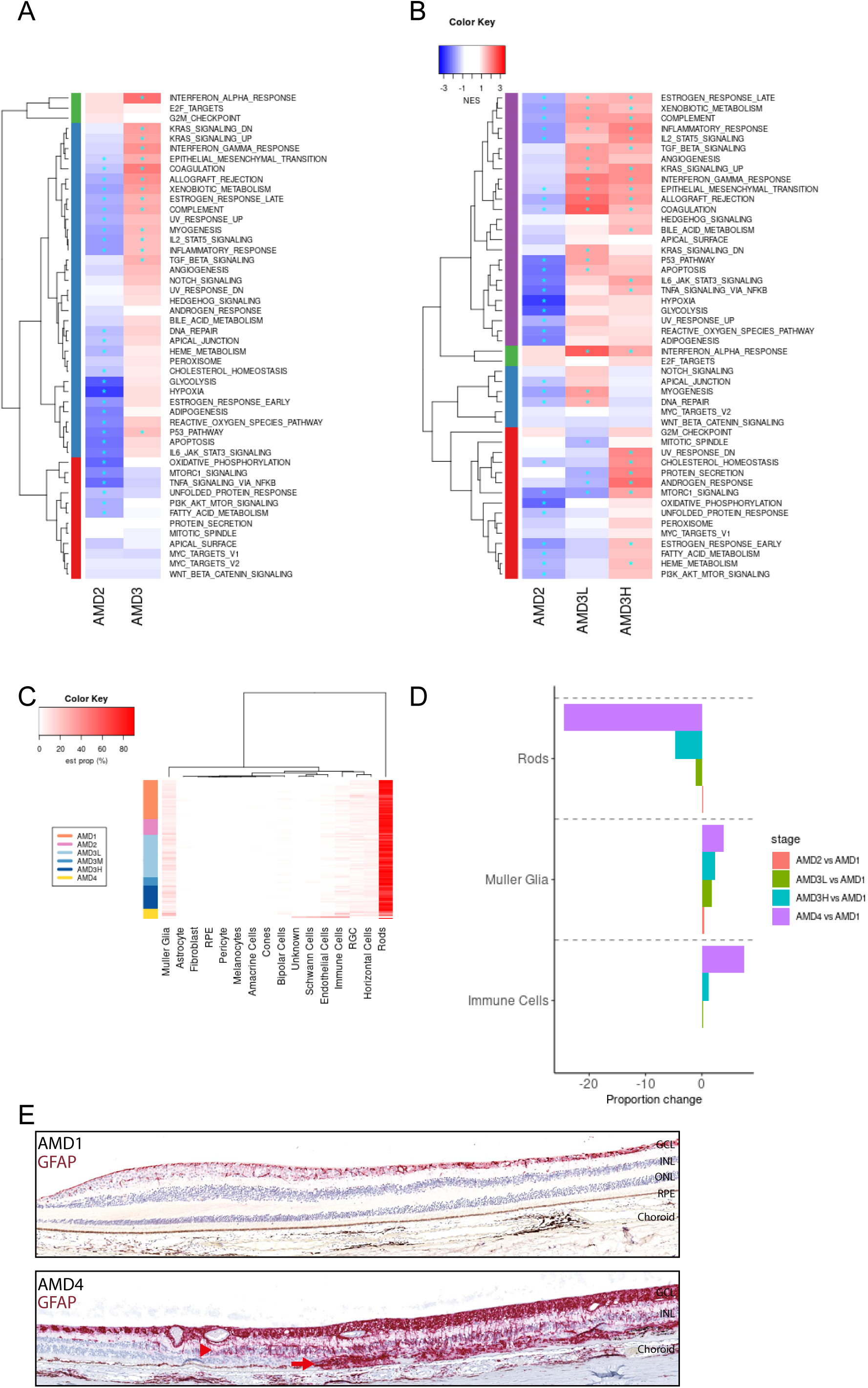
In macular neural retina bulk RNAseq, major pathway and cellular changes occur in advanced stages of AMD. **A**. Heatmap of normalized enrichment scores (NES) of GSEA Hallmark gene sets from macular neural retina bulk RNAseq data comparing each AMD stage with AMD1. Upregulated pathways are indicated in red and downregulated ones in blue. An asterisk (*) indicates any FDR < 0.05. The sample numbers for AMD1, 2, 3L, 3H, and 4 are 26, 10, 15 and 7, respectively. We also included all AMD3 samples (N = 27) from MGS4 grading in the heatmap in comparison to the AMD3L and AMD3H samples from MGS9 grading. AMD3M (N=6) was excluded from the comparison due to smaller sample size. **B.** Cell proportions in the macular neural retina across AMD grades as inferred by MuSiC. Rod photoreceptors comprise the bulk of each sample, followed by Müller glia and horizontal cells. The term “est prop (%)” refers to the estimated proportion of a given cell type. **C.** Average cell proportion of selected cell types in each AMD stage relative to AMD1. The proportion of rod photoreceptors is reduced in AMD4, whereas the proportions of Müller glia and immune cells increased in AMD4. **D.** Representative images of IHC staining of GFAP (red), an astrocyte and Müller glial activation marker, with macular cross-sections from representative AMD1 and AMD4 donors. Scale bar = 100 µm. The arrowhead shows increasing GFAP staining in the transition zone, defined as an area with RPE abnormality and PR thinning. The arrow shows GFAP signaling extending across the atrophic area, defined as absence of RPE and PR loss.

### 3.4 Validation of molecular signatures associated with a high risk of progression to late stages of AMD

Stratifying AMD3 using the MGS9 grading system allowed us to identify gene signatures associated with intermediate AMD samples that were at a high risk of progression to end-stage disease. We next aimed to identify specific genes that could serve as markers of progression, by evaluating top ranked genes that expressed higher in the macular RPE/choroid of AMD3H samples than in AMD3L samples, compared to AMD1 (**Figure 4A**). Out of the top 150 differentially expressed genes, 7 genes were selected for further confirmation based on their cell-type/tissue-specific expression and potential functional relevance to AMD pathobiology (**Table 1**, **Figure 4B**). These markers represent activation of complement signaling, macrophage markers, and astrocyte/Müller glia markers, based on assessment of snRNA-seq data from the macular RPE/choroid and neural retina (**Supplemental Figure 4**).

**Figure 4.**
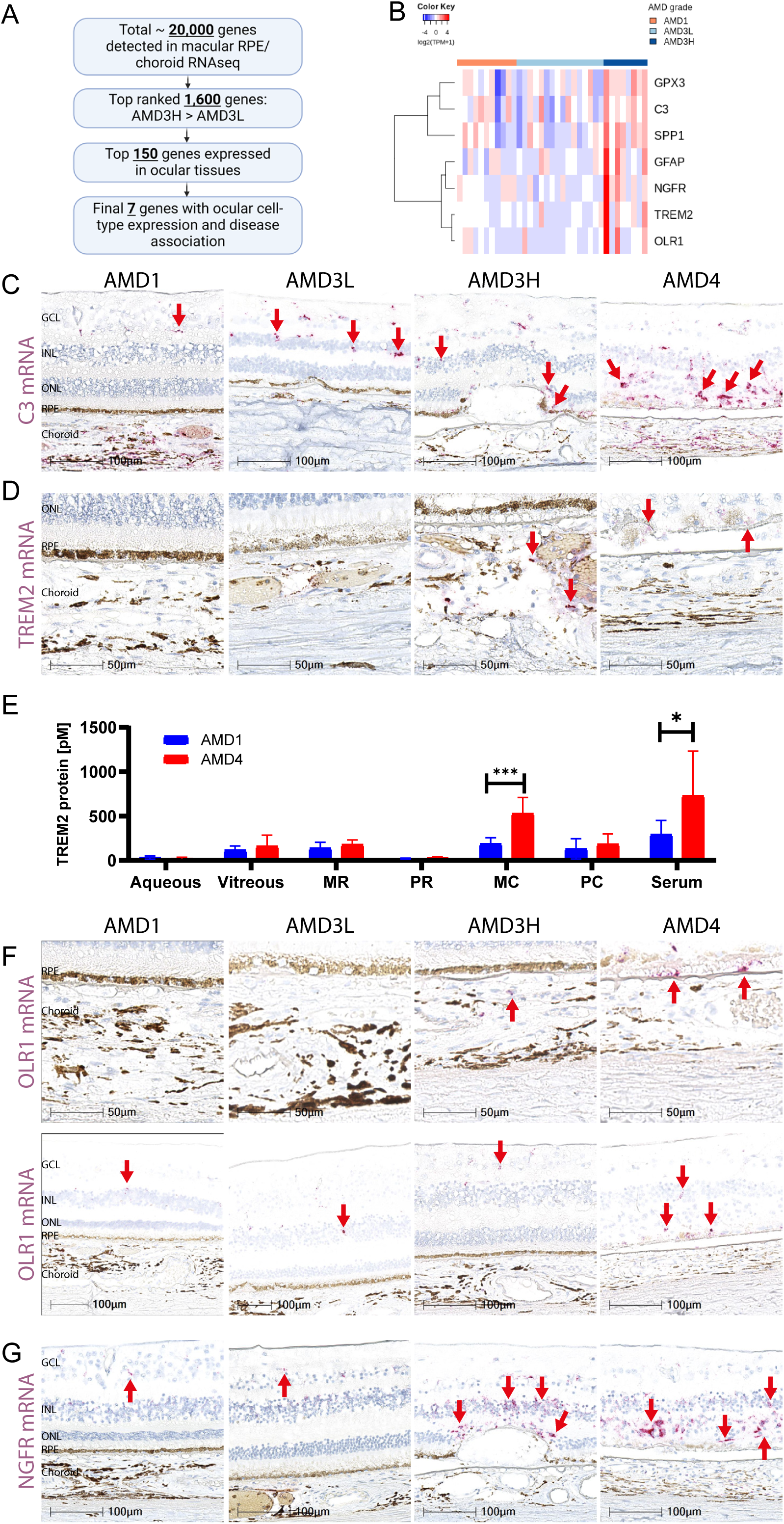
Molecular signatures indicating a high risk of progression to the late stage of AMD. **A**. Flowchart describing selection of gene markers associated with a high-risk of progression to late AMD for validation, based on bulk RNAseq of the macular RPE/choroid. **B.** A heatmap showing high-risk marker gene expression in the macular RPE/choroid in samples of AMD1, AMD3L, and AMD3H. The levels of each mRNA are expressed as log 2 (TPM+1), where TPM stands for transcript per million, red indicates positive value and blue negative. **C**. Representative RNAscope images of *C3* mRNA in AMD1, AMD3L, AMD3H, and AMD4/GA macula sections, with arrows indicating *C3* mRNA^+^ signals. *C3* increases in the RPE-adjacent region with disease severity. **D.** Representative RNAscope images of TREM2 mRNA in AMD1, AMD3L, AMD3H, and AMD4/GA macula sections, with arrows indicating TREM2 mRNA+ signals. *TREM2* mRNA becomes apparent in the choroid at AMD3H and persists in AMD4. **E.** ELISA measurement of TREM2 protein in AMD1 (n = 8 samples) and AMD4 (n = 6-8) in serum, macular and peripheral RPE/choroid tissue and neuroretina, vitreous, and aqueous humor. Donor information is in **Supplemental Table 7**. The pM label indicates picomolar concentrations, with p value determined by unpaired Student’s t-test (* p < 0.05, *** P <0.001). **F**. Representative RNAscope images of *OLR1* mRNA in AMD1, AMD3L, AMD3H, and AMD4/GA macula sections, with arrows indicating *OLR1* mRNA+ signals. *OLR1* increases in the RPE-adjacent region with disease severity. **G**. Representative RNAscope images of NGFR mRNA in AMD1, AMD3L, AMD3H, and AMD4/GA macula sections, with arrows indicating *NGFR* mRNA+ signals. Elevated *NGFR* signal is observed in AMD3H, proximal to the RPE.

**Table 1.**
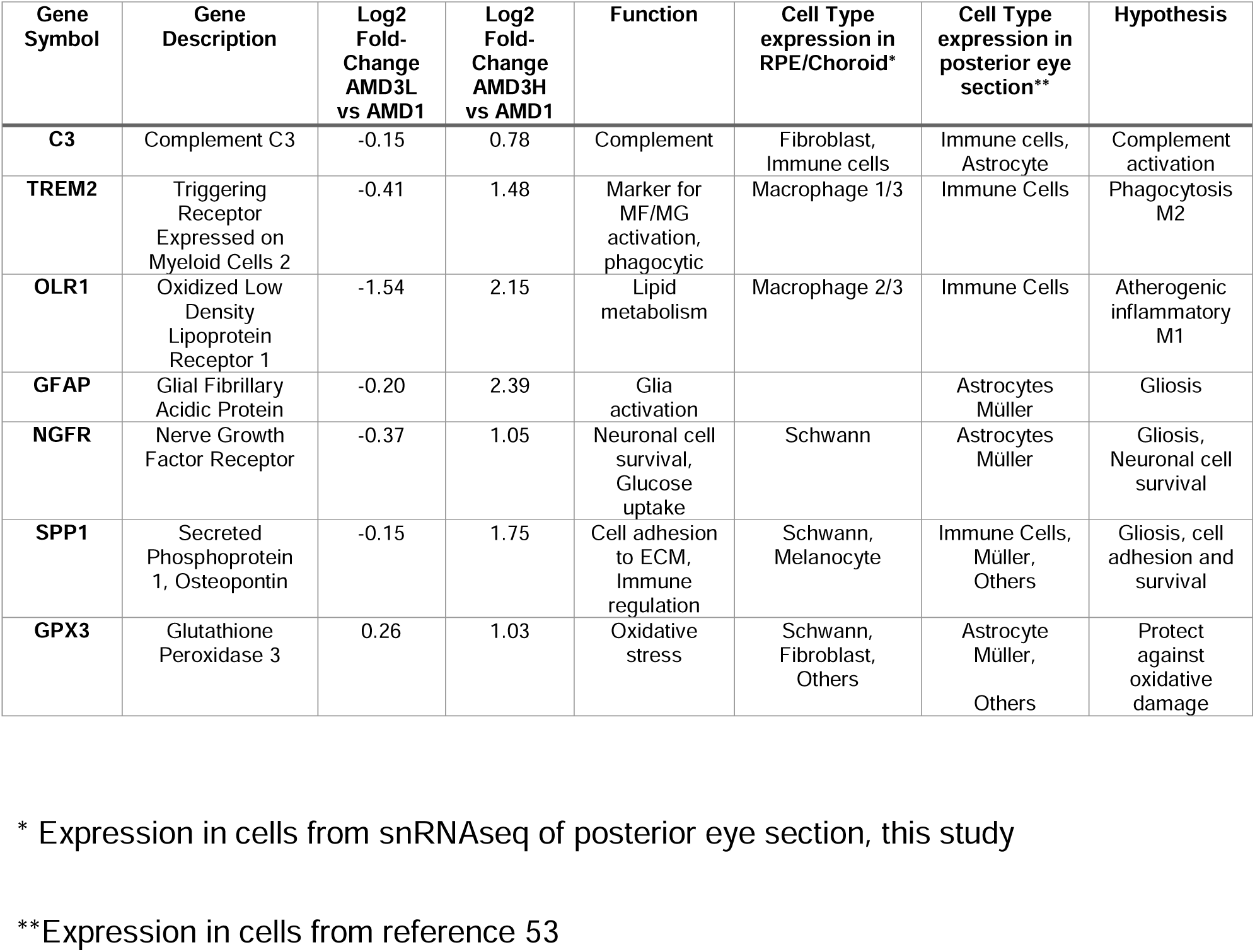
Summary of intermediate AMD high risk genes selected for validation.

We could not directly apply the MGS9 system to eyes collected for histological validation of our selected genes, as these eyes were collected as fixed globes without macular fundus imaging. By removing the neural retina to accurately grade the eye according to MGS9, the opportunity to perform histology on the same globe is lost. Therefore, we graded histology eyes according to the Sarks grading system ^40,42^ starting with a four-stage severity scale of AMD based on H&E histopathological features as previously described ^26^. In order to correlate with the AMD3L/M/H substratification from dissected post-mortem eyes, AMD3 histology samples were further divided into three sub-groups (**Figure S2;** refer to **Supplemental Table 6A and 6B** for donor information and sample allocation) based on the severity of the AMD histopathology as per Starks’ AMD pathological characterization ^43^. AMD3L macular sections exhibited RPE rounding and disrupted cobblestone morphology, melanin sloughing, and thin patchy basal lamina deposits (BLmD) with an accumulation of sub-RPE material thickness less than 7 µm between RPE and BrM. AMD3M features include the abnormalities seen in AMD3L, plus disorganized outer segments, loss of the outer nuclear layer (ONL), and thin continuous BLmD accumulation of sub-RPE material thickness that was equal or less than 7 µm in thickness. In addition to AMD3M characteristics, AMD3H eyes displayed thick continuous BLmD accumulation of sub-RPE material thickness equal or greater than 7 µm, extending over 250 µm in length underneath the RPE cells, sparse RPE thinning, extensive RPE hypertrophy, and BrM hyalinization.

Complement activation has been strongly implicated in AMD pathogenesis ^4,5,44^ and intravitreal injection of the C3 inhibitor, Pegcetacoplan (SYFOVRE®), is the first FDA-approved treatment for GA ^45,46^. Interestingly, the central complement component *C3* appeared as one of the high-risk genes for AMD progression from our macular RPE/choroid gene profiling. Consistent with previous reports that the RPE/choroid complex constitutes the main source of complement C3 expression and activation in AMD ^26,47^, we observed numerous *C3+* cells in the GA lesion areas (**Figure 4C**). In healthy eyes, C3 mRNA^+^ cells which appear microglia-like based on morphology are restricted to the inner retina, while the PR/RPE/choriocapillaris area is devoid of *C3* mRNA. Induction of *C3* mRNA signals in the outer retinal region, often near or within the RPE layer, occurs in AMD3H. Co-labelling of C3 mRNA with IBA1 revealed presence of dual positive cells in the ONL/RPE complex (**Supplemental Figure S5**). C3 mRNA+/IBA1+ cells closely associated with the RPE also appear pigmented, suggesting that macrophages and activated microglia may both contribute to intraocular complement activation via increasing local complement gene expression and to RPE phagocytosis (**Supplemental Figure S5**). Fibroblasts could also be additional sources of complement in the choroidal space, as they also show *C3* expression at baseline in snRNAseq data. (**Supplemental Figure S3**).

We identified the myeloid phagocytic membrane receptor *TREM2* as an AMD high-risk gene. As a major regulator of monocellular phagocytes’ phagocytosis, migration, and immunomodulatory state, TREM2 is a key risk factor for other age-associated diseases with similar pathophysiology to AMD. Interestingly, decreased expression, functional impairment, and genetic variants of TREM2 have been found associated with neurodegenerative diseases such as Alzheimer’s disease (AD), Parkinson’s disease, and frontotemporal dementia ^48,49^ and several clinical trials are currently exploring the potential of TREM2 agonism as therapeutic intervention in AD ^50^. Similar to neurodegenerative diseases, a role for TREM2 in promoting neuroprotective functions of microglia in AMD has been recently proposed ^51,52^. In our donor samples, we only observed a limited number of *TREM2* mRNA^+^ cells in the back of the eye, specifically near the choriocapillaris in AMD3H and around the GA lesions in AMD4 (**Figure 4D**). These findings were confirmed by ELISA quantification of total TREM2 protein showing enrichment in the macular RPE/choroid compared with other ocular tissues (**Figure 4E**). Importantly, TREM2 protein levels in both the macular RPE/choroid and serum were significantly increased in GA/AMD4, suggesting that *TREM2+* myeloid cells, likely macrophages, are recruited to phagocytose dying RPE and photoreceptors. This contrasts with prior reports of decreased TREM2 protein levels in macular retinas of dry AMD donors ^52^ which may be explained by the subretinal translocation and enrichment of TREM2+ microglia at the diseased/atrophic sites challenging retrieval upon neural retina dissection from the posterior eyecup. Another high-risk gene, *OLR1* (Oxidized Low Density Lipoprotein Receptor 1), known to mediate the polarization and pro-inflammatory status of macrophages, was found to be enriched in AMD3H choroid and in the lipid rich soft drusen areas of AMD4 samples (**Figure 4F**). A prior snRNAseq study ^53^ suggests that *OLR1+* macrophages may represent a subpopulation of macrophages distinct from *TREM2^+^* (**Supplemental Figure S6**), suggestive of interplay between multiple types of macrophages in AMD progression.

Markers of Müller/astrocyte glia activation in AMD have been previously reported to be upregulated in AMD ^24,54,55^. Accordingly, multiple AMD high-risk genes are preferentially expressed in Müller glia and astrocytes including *GFAP*, *NGFR*, and *GPX3* (**Table 1, Supplemental Figure S4**). *NGFR*, the nerve growth factor receptor, appears to be more specifically expressed in astrocytes in the neural retina and Schwann cells in the RPE/choroid. While only a few copies of *NGFR* mRNA are detected in the inner, but not outer, retina from AMD1 to AMD3L, it is abundantly present around the RPE atrophic area in AMD3H and AMD4 (**Figure 4G**), indicating an active involvement of NGFR during RPE/PR degeneration. *SPP1* (Secreted Phosphoprotein 1) and *GPX3* (Glutathione Peroxidase 3) are broadly expressed in all ocular cell-types with relatively higher expression in astrocyte, Müller glia, immune cells, and Schwann cells (**Supplemental Figure S4**). RNAscope of *SPP1* and *GPX3* confirms their widespread expression as well as their upregulation in AMD3H/4 (**Supplemental Figure S7).**

In summary, we have identified and confirmed several molecular and cellular high-risk markers during the progression to the late/advanced stage of AMD as summarized in **Figure 5**. Complement *C3* is upregulated likely in fibroblasts, macrophages, and other cell types in AMD3H and AMD4. *TREM2^+^* and *OLR1^+^* macrophages are present around the BrM/choroid in AMD3H, and then in the GA lesion area of AMD4. Müller/astrocyte expressed gene/proteins (*NGFR*, GFAP) extend from the inner retina to outer retina and sub-retinal/sub-RPE space in AMD4. The association of some of these markers with AMD has been previously reported ^19,26,51,52,56^, while others will need further independent confirmation.

**Figure 5.**
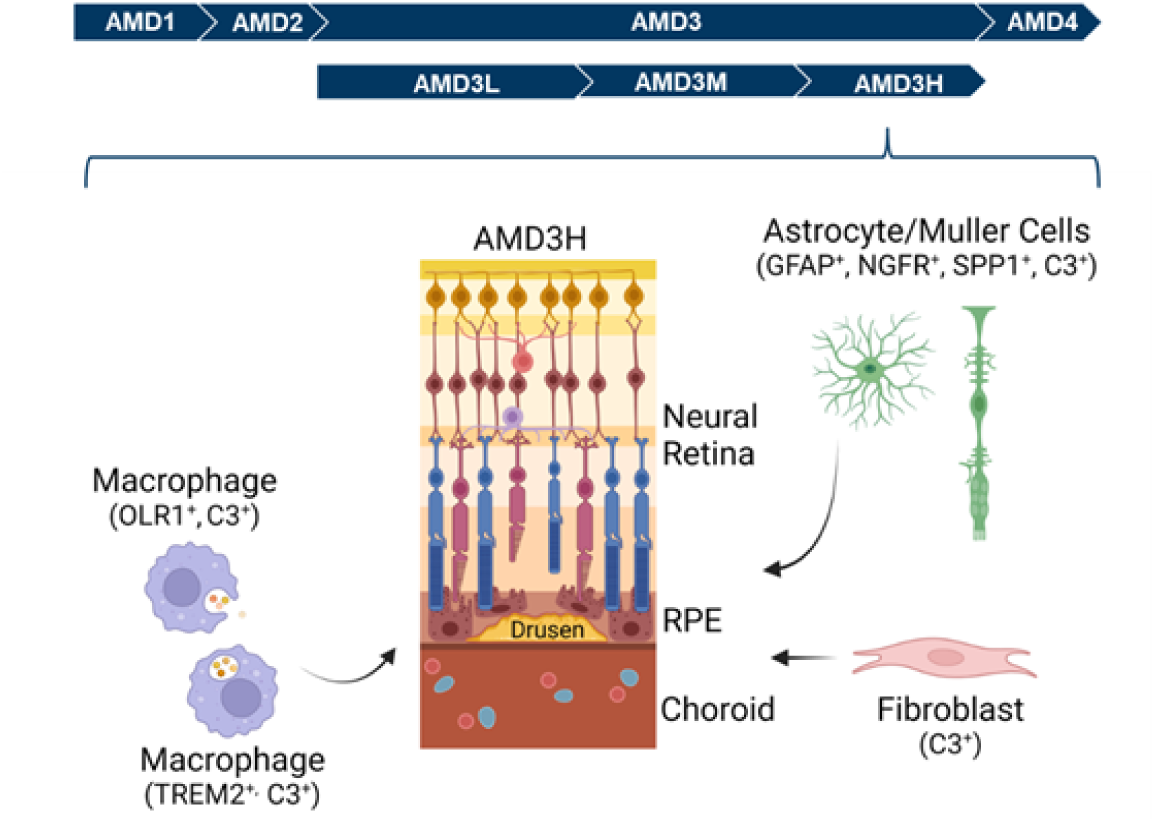
Schematic presentation of high risk cellular and molecular signatures observed during progression of AMD from intermediate to geographic atrophy. In AMD3H, complement *C3* can be induced in multiple neuronal and choroidal cell-types including fibroblasts, macrophages, and Müller/astrocytes. Macrophages expressing either *TREM2* or *OLR1* are observed near the RPE/choriocapillaris complex in AMD3H, possibly originating from the choroid and/or from the systemic circulation, and later in AMD3H/4 outer retina. Activated Müller/astrocytes cells expressing *NGFR* and GFAP expand from the inner retina to outer retina and RPE in AMD3H. Activation of complement, macrophages, and Müller/astrocytes may signal the onset of the end stage AMD. Graphic was created with Biorender (biorender.com).

## 4. Conclusion

By leveraging the advantages of the more detailed MGS9 grading scale, we are able to further divide the complex and the heterogenous intermediate AMD (AMD3) stage for postmortem eyes into 3 subgroups. These subgroups - AMD3L, AMD3M and AMD3H – are based on the binned risk of disease progression to late AMD. This substratification allowed us to observe that opposing transcriptional changes ascribed to RPE and immune cells occur between the AMD3L and AMD3H stages in the macular RPE/choroid. In the macular neural retina, we found that most biological pathways were downregulated in the AMD2 stage but upregulated in AMD3L and AMD3H, with significantly increased expression at the individual gene level in AMD4. The higher expression of RPE genes and lower expression of retinal genes in AMD2 and AMD3L, as assessed by single-cell mediated deconvolution of bulk RNA sequencing data, may represent a compensatory response designed to minimize the pathogenic effect in the early stage of the disease. However, by stages AMD3H and AMD4, the inflammatory response and gliosis become dominant. Importantly, we identified and confirmed distinct molecular and cellular signatures indicating a high risk of progression to late AMD. These include activation of complement (C3), macrophage-specific genes (TREM2, OLR1), and alterations in Müller/astrocyte glia (evidenced by GFAP, NGFR). The identification and ocular localization of these high-risk biomarkers, as demonstrated in this study, hold significant value for improving AMD sub-stratification of donor tissues such as retinal sections via histological analysis. More importantly, they may provide novel insights into the key biological processes underlying the conversion to atrophic degeneration during AMD progression and identify novel therapeutic targets. However, it remains critical to integrate these differential expression findings with functional analysis and mechanistic biological studies to establish clear causative pathways and infer proper directionality for therapeutic treatments.

## Abbreviations

AMD: age-related macular degeneration
BrM: Bruch’s membrane
CC: choriocapillaris
GA: geographic atrophy
GCL: ganglion cell layer
INL: inner nuclear layer
MGS: Minnesota grading system
MGS4: 4-step MGS
MGS9: 9-step MGS
nAMD: neovascular AMD
NES: Normalized Enrichment Score
ONL: outer nuclear layer
PR: photoreceptor
RPE: retinal pigment epithelium

## Author contributions

C.-L. H., J.D., M. T., O. D., M. C., N. B., N. V., L. F., X. W., J. Y., T. R. V., G. K., T. W. O.– Investigation; Y.Q., J.D., J. A., S. P., C. W. W., S.-M. L. – Conceptualization; C.-L. H., S.-M. L.–Formal analysis; C.-L. H., Y. W., S. J., T. W. O.– Data Curation; C.-L. H., Y. Q., J. D., M. C., N. B., L. F., M. T., J. A., M. S.-G., G. P., S. P., C. W. W., C. G., T. W. O., S.-M. L. – Writing - Review & Editing; C.-L. H., Y. Q., J. D., M. T., X. W., S.-M. L.– Visualization; Y.Q., C. W. W. – Resources; M. S.-G., G. P., S. P., C. W. W., C. G.- Supervision; C.W.W., M.S.-G., S.-M. L.– Writing - Original Draft and Project Administration

## Funding

This work was supported by Novartis Biomedical Research.

## Declaration of competing interests/financial relationships

The following authors were Novartis employees at the time of this research study and may hold stock in the company: Chia-Ling Huang (E), Yubin Qiu (E), John T. Demirs (E), Xiaoqiu Wu (E), Michael Twarog (E), Omar Delgado (E), Maura Crowley (E), Natasha Buchanan (E), Nhi Vo (E), Lin Fan (E), Yanqun Wang (E), Junzheng Yang (E), Thomas R. Vollmer (E), Garret Klokman (E), Sandra Jose (E), Jorge Aranda (E), Ganesh Prasanna (E), Stephen Poor (E), Cynthia Grosskreutz (E), Christopher W. Wilson (E), Magali Saint-Geniez (E), Sha-Mei Liao (E). Other relationships: Timothy Olsen (C).

## Acknowledgements

The authors thank Thaddeus Dryja, Chen Yu, and Sarah Richards for their comments and suggestions. We thank Nicholas Sprehe, David Ammar, Jason Woody and the Lions World Vision Institute for procurement and dissection of human donor eyes. The authors are thankful to the donors and their families for their valuable gift to research.

## SUPPLEMENTAL FIGURE LEGENDS

**Supplemental Figure S1.**
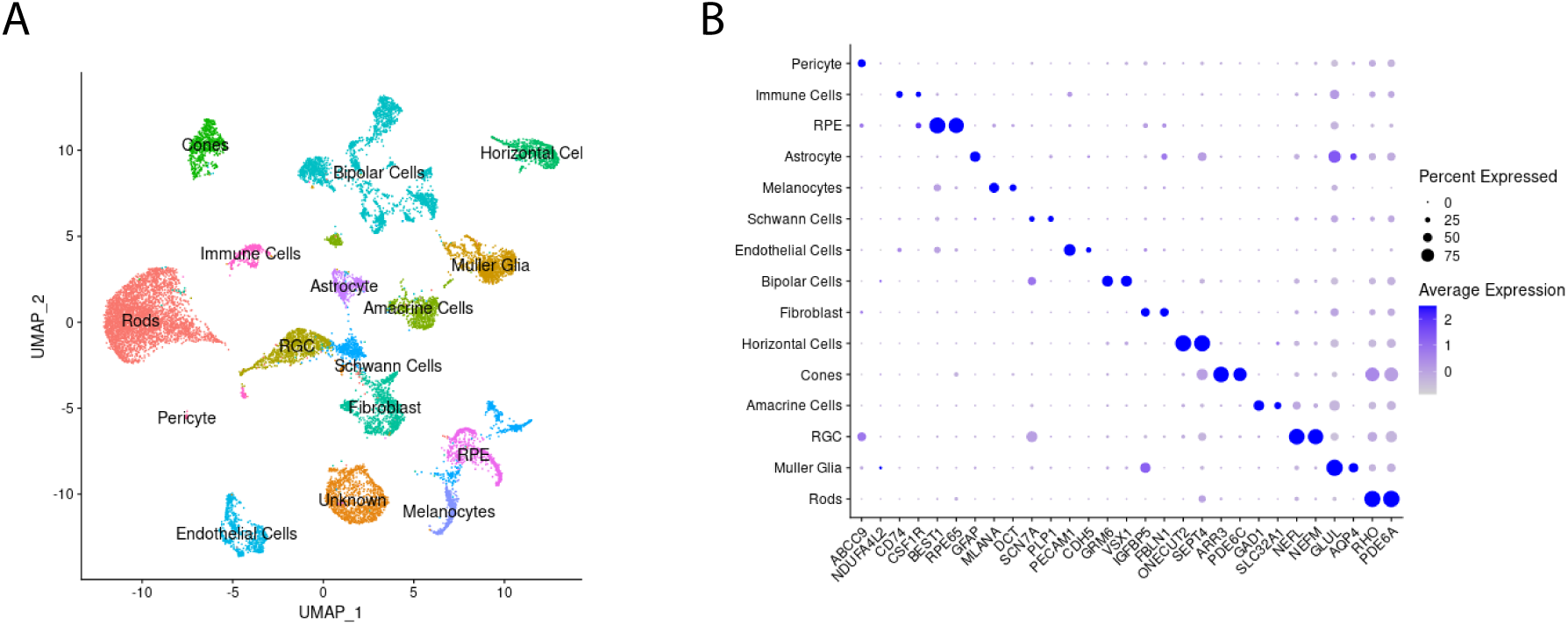
Single nuclei RNA sequencing of human retina/RPE/choroid from frozen tissue. **A**. UMAP visualization of cell clusters identified from human retina/RPE/Choroid snRNAseq from frozen sections. **B**. Expression of a subset of selected cell-type marker genes used for cluster annotation.

**Supplemental Figure S2.**
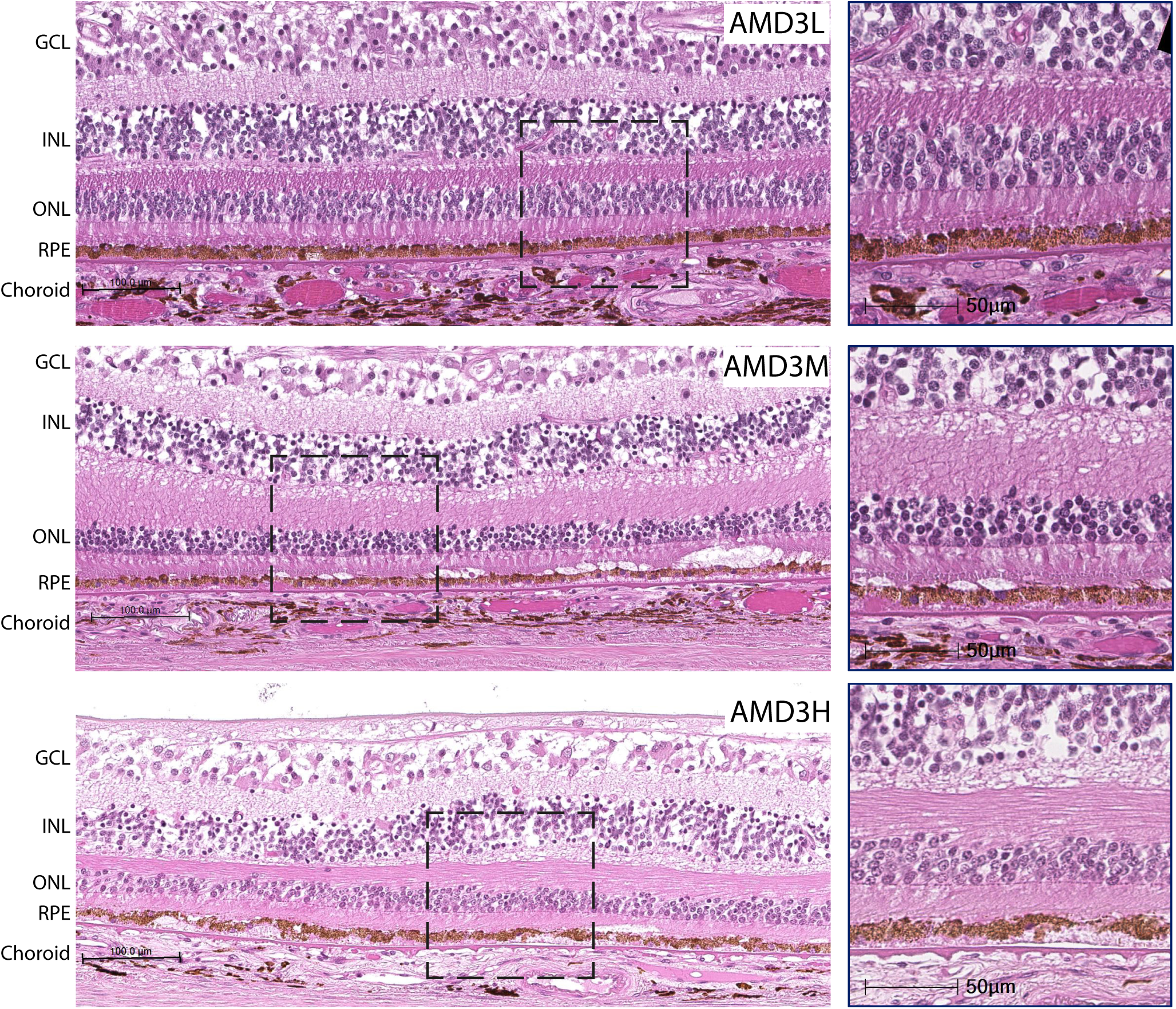
Representative haematoxylin and eosin (H&E) stained images of AMD3L, 3M, 3H cross-sections. Images on the right are magnified areas from the corresponding left images. AMD3L displays RPE rounding (cobblestone), melanin sloughing, and thin patchy basal lamina deposit (BLmD) accumulation of sub-RPE material thickness less than 7 µm. AMD3M shows disorganized outer segments, loss of the outer nuclear layer (ONL), and BLmD thickness equal or less than 7 µm in addition to AMD3L pathology. AMD3H displays BLmD thickness greater than 7 µm, extending > 250 µm, and BrM hyalinization (glassy appearance), in addition to all AMD3M and AMD3L pathology. Scale bar = 100 µm.

**Supplemental Figure S3.**
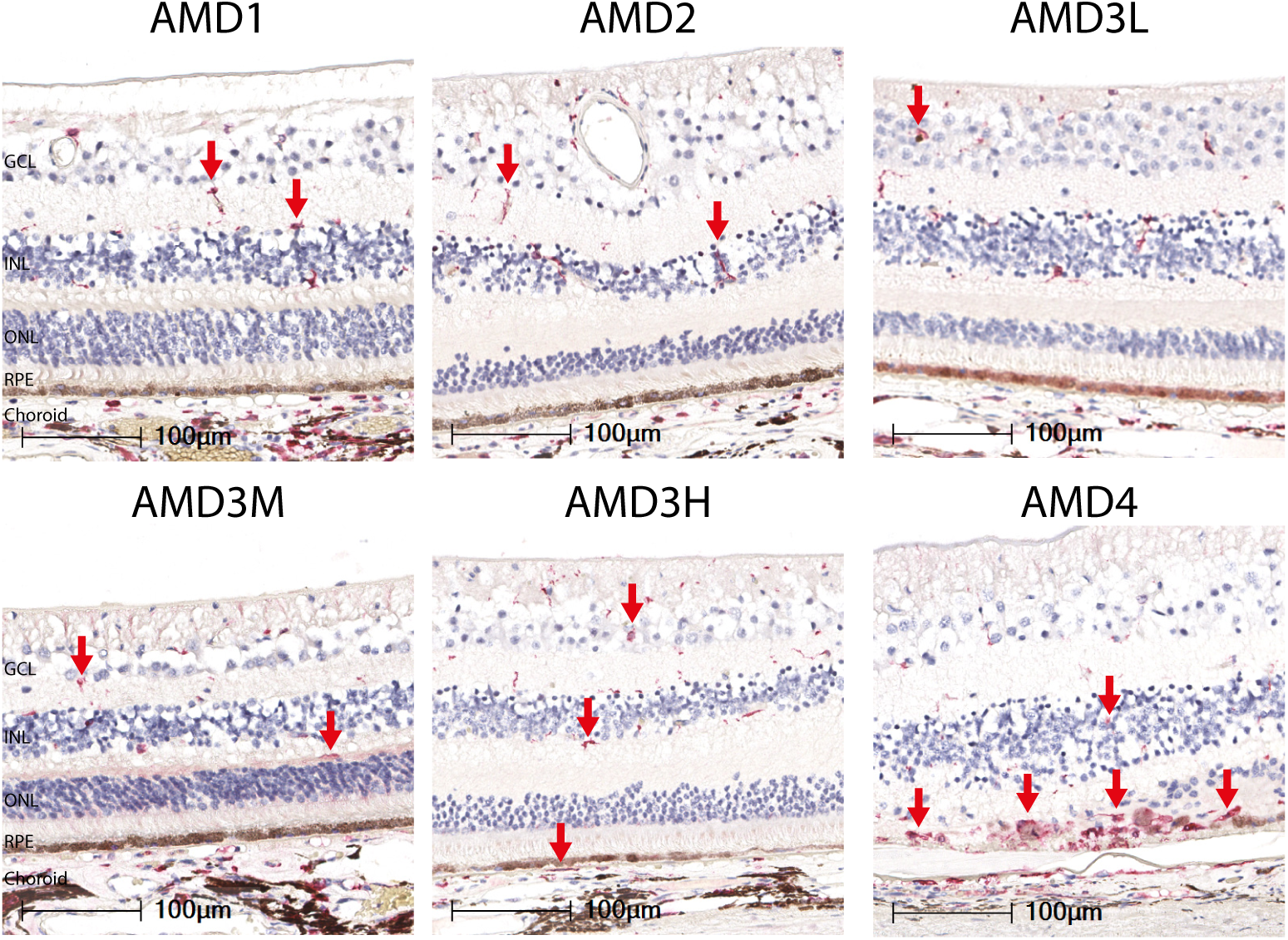
Representative images of IBA1 staining of retinal sections across AMD grades. In AMD1, IBA1 positive cells primarily localize to the inner retina, where they exhibit a ramified structure and do not extend beyond the OPL layer. As AMD progresses through AMD2 and AMD3 sub-stages, IBA1 positivity increases while still largely retaining localization in the inner retina. In AMD4, a pronounced increase in IBA1 positivity is observed in the outer retina, primarily in the atrophic lesion are, with cells exhibiting amoeboid morphology. This, coupled with a qualitative reduction of IBA1+ cells in the inner retina, suggests possible migration from the inner retina, or from the choroid through the damaged RPE and Bruch’s membrane.

**Supplemental Figure S4.**
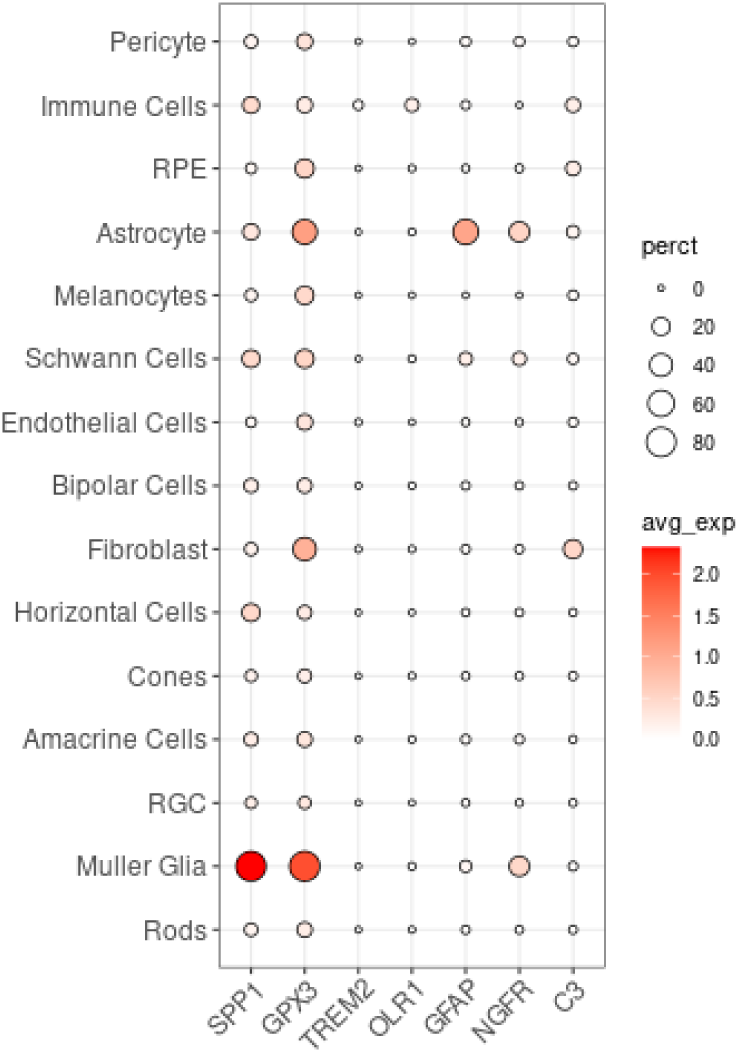
Dot plot of cell-type specific gene expression for the seven selected AMD high-risk progression marker genes in human retina/RPE/choroid. Fibroblasts predominantly express C3, immune cells show C3, OLR1 and TREM2 expression, and GFAP, NGFR, SPP1 and GPX2 are expressed in Muller glia and astrocytes.

**Supplemental Figure S5.**
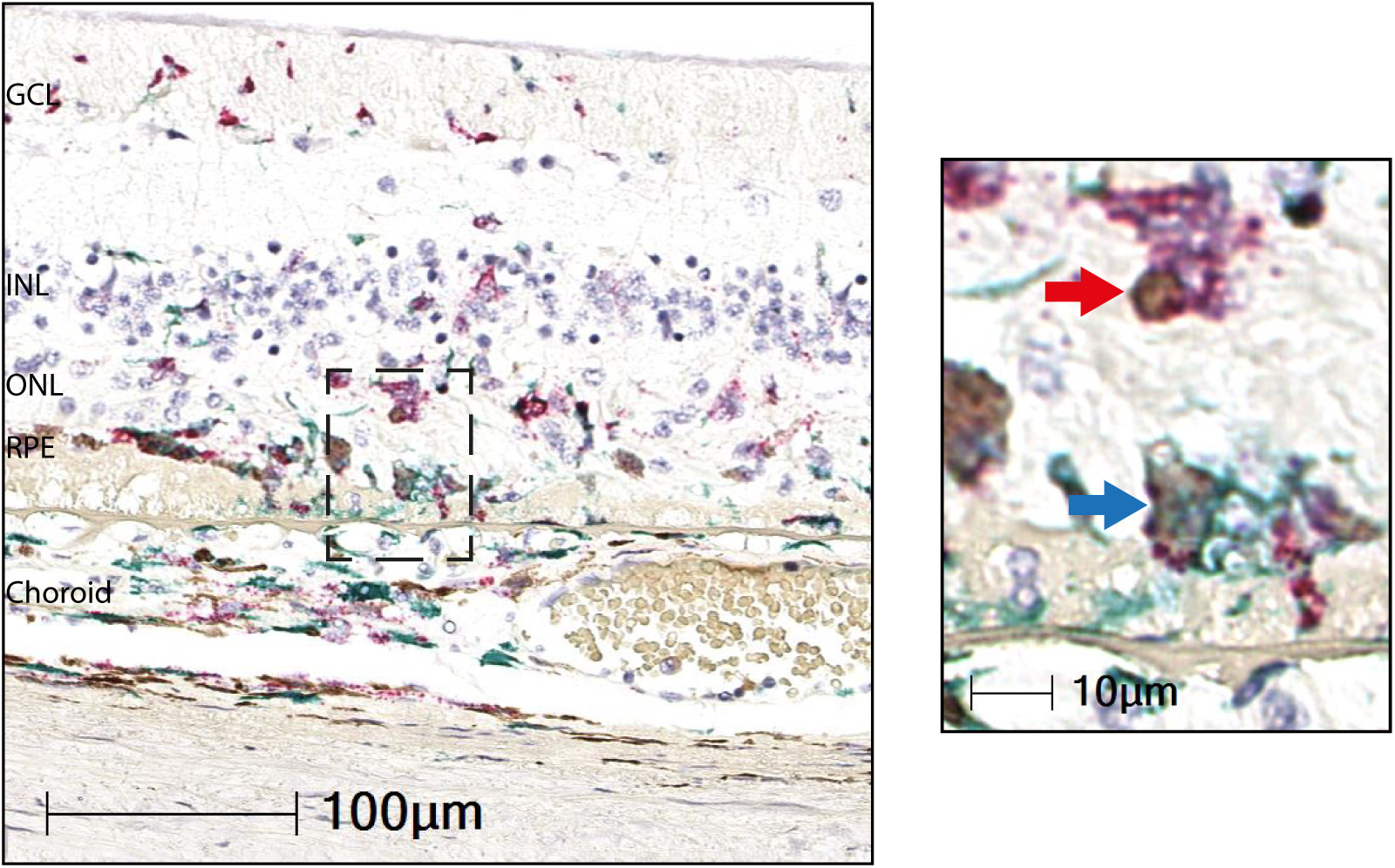
Costaining of *C3* mRNA (red) and IBA1 protein (green) in an atrophic GA/AMD4 section. In the inset, the green arrow indicates an IBA1 and *C3* mRNA double positive cell. The red arrow indicates a *C3* positive but IBA1 negative cell.

**Supplemental Figure S6.**
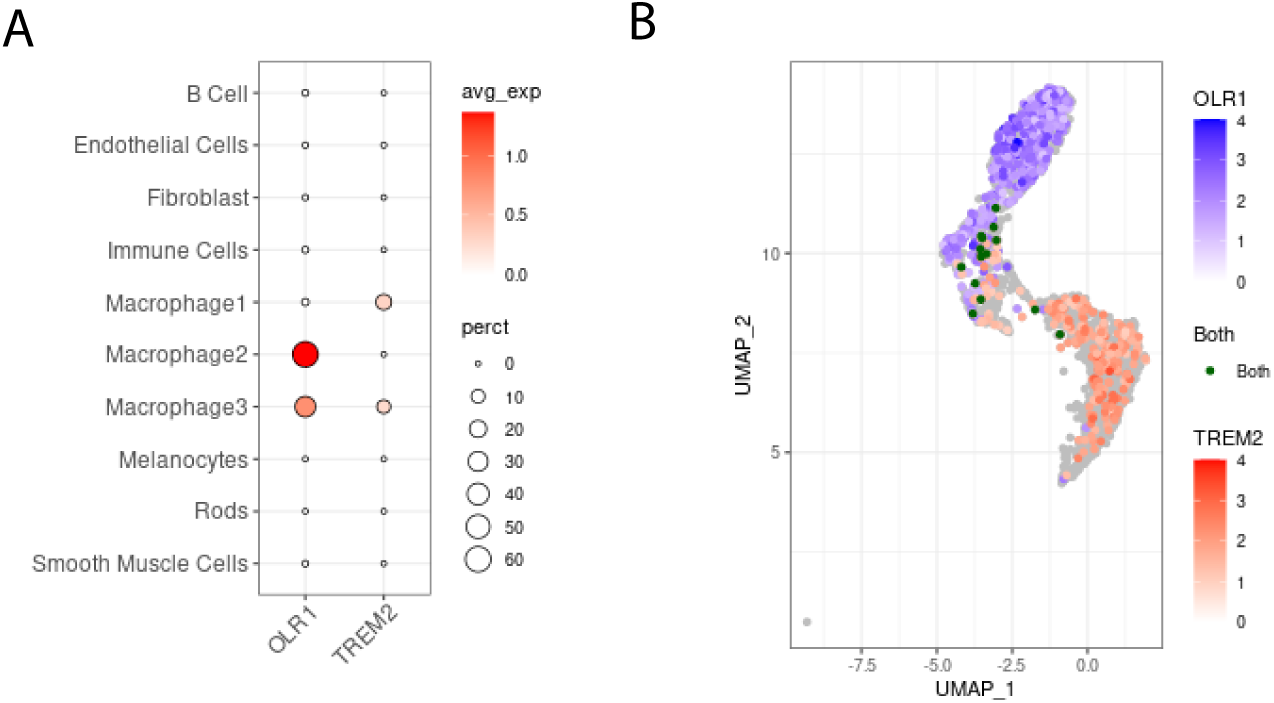
*TREM2* and *OLR1* are preferentially expressed in different types of choroidal macrophages in published single cell RNA-seq from human RPE/choroid samples ^53,57^. **A**. A dot plot of *TREM2* and *OLR1* expression in all detected RPE/choroid cell types. Both*TREM2* and *OLR2* are preferentially expressed in choroidal macrophages. Of the three clusters of macrophages, *TREM2* is expressed in macrophage 1 and 3 while *OLR1* is expressed in macrophage 2 and 3. **B**. Dual genes plot of all 3 macrophage populations expressing *TREM2* (green dots) versus *OLR1* (red dots). There are very few macrophages (yellow dots) expressing both *TREM2* and *OLR1*.

**Supplemental Figure S7.**
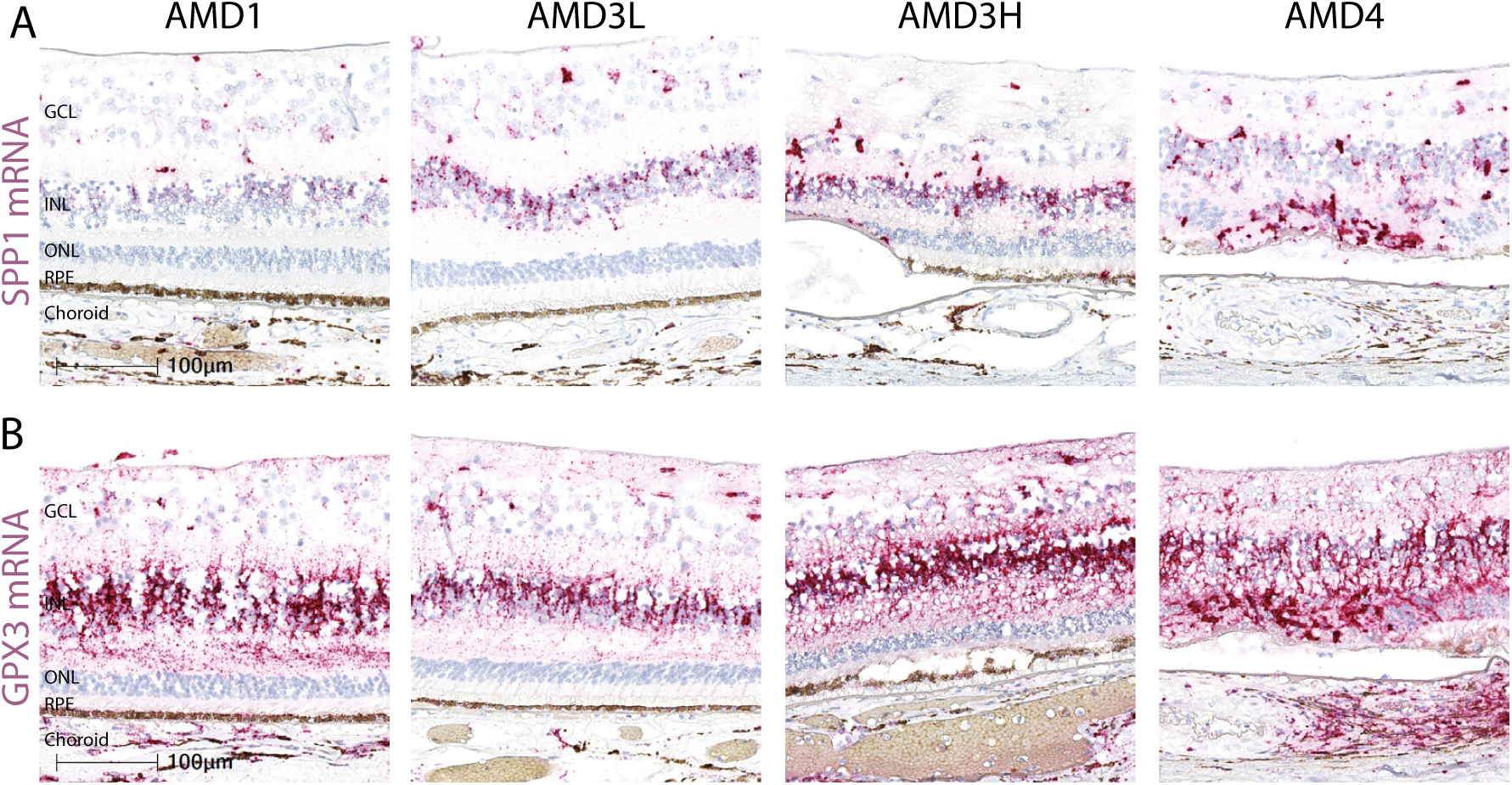
RNAscope of *SPP1* and *GPX3* in the neural retina. Representative RNAscope images of *SPP1* (A) or *GPX3* (B) mRNA in AMD1, AMD3L, AMD3H, and AMD4/GA macula sections. There are many cells expressing either *SPP1* or *GPX3* mRNA in the retina cross-sections. Consistent with RNA-seq data, there appear to be more *SPP1* and *GPX1* mRNA copies in AMD3H and AMD4 than in AMD1 and AMD3L.

## SUPPLEMENTAL TABLES

**Supplemental Table 1.**
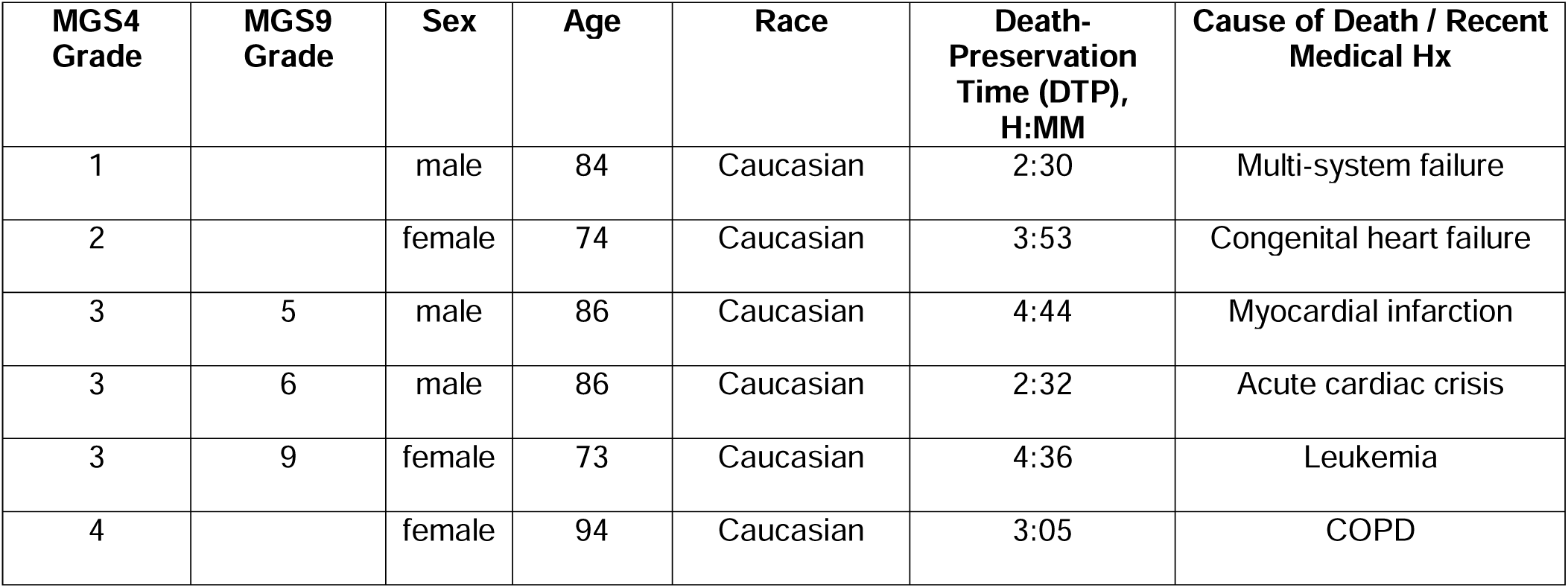
Donor demographics for postmortem eyes in Figure 1B.

| **MGS4**<br>**Grade** | **MGS9**<br>**Grade** | **Sex** | **Age** | **Race** | **Death-**<br>**Preservation**<br>**Time (DTP),**<br>**H:MM** | **Cause of Death / Recent**<br>**Medical Hx** |
| --- | --- | --- | --- | --- | --- | --- |
| 1 |  | male | 84 | Caucasian | 2:30 | Multi-system failure |
| 2 |  | female | 74 | Caucasian | 3:53 | Congenital heart failure |
| 3 | 5 | male | 86 | Caucasian | 4:44 | Myocardial infarction |
| 3 | 6 | male | 86 | Caucasian | 2:32 | Acute cardiac crisis |
| 3 | 9 | female | 73 | Caucasian | 4:36 | Leukemia |
| 4 |  | female | 94 | Caucasian | 3:05 | COPD |

**Supplemental Table 2A.**
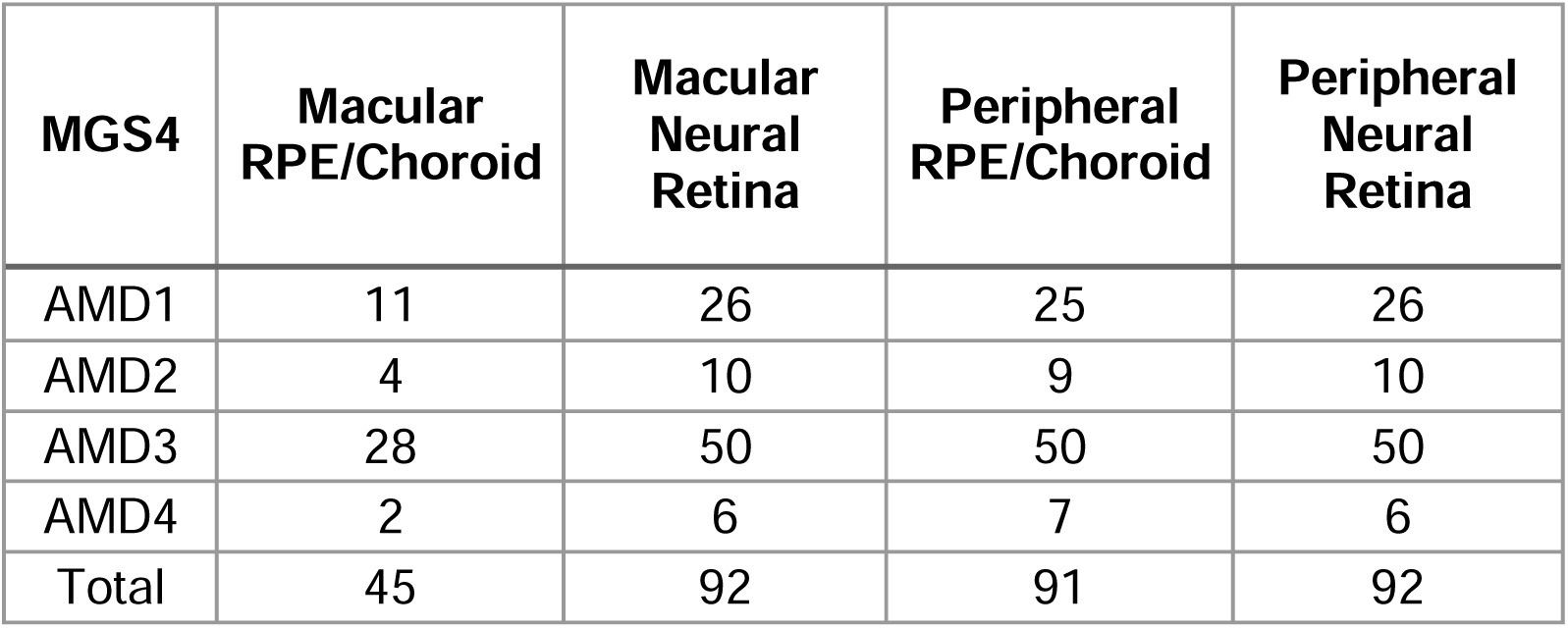
Postmortem eyes used for bulk RNAseq, grouped by MGS4.

| **MGS4** | **Macular**<br>**RPE/Choroid** | **Macular**<br>**Neural**<br>**Retina** | **Peripheral**<br>**RPE/Choroid** | **Peripheral**<br>**Neural**<br>**Retina** |
| --- | --- | --- | --- | --- |
| AMD1 | 11 | 26 | 25 | 26 |
| AMD2 | 4 | 10 | 9 | 10 |
| AMD3 | 28 | 50 | 50 | 50 |
| AMD4 | 2 | 6 | 7 | 6 |
| Total | 45 | 92 | 91 | 92 |

**Supplemental Table 2B.** Postmortem eyes used for bulk RNAseq, grouped by MGS9.

| **MGS9** | **Macular**<br>**RPE/Choroid** | **Macular**<br>**Neural**<br>**Retina** | **Peripheral**<br>**RPE/Choroid** | **Peripheral**<br>**Neural**<br>**Retina** |
| --- | --- | --- | --- | --- |
| AMD1 | 11 | 26 | 25 | 26 |
| AMD2 | 4 | 11 | 10 | 11 |
| AMD3L | 16 | 27 | 26 | 27 |
| AMD3M | 4 | 6 | 7 | 6 |
| AMD3H | 8 | 15 | 15 | 15 |
| AMD4 | 2 | 6 | 7 | 6 |
| Total | 45 | 91 | 90 | 91 |

**Supplemental Table 3.** Donor demographics and regions used from postmortem eyes in bulk RNAseq.

| Sample ID | Eye | MGS4 | MGS9 | MGS (3 bins) | Age | Sex | Race | DTP (Hours) | Cause of Death / Recent Medical Hx | MC | PC | MR | PR |
| --- | --- | --- | --- | --- | --- | --- | --- | --- | --- | --- | --- | --- | --- |
| Sample1 | OD | AMD3 | 2 | NA | 92 | male | caucasian | 3:29 | Pneumonia | X | X | X | X |
| Sample2 | OS | AMD3 | 4 | AMD3L | 90 | female | caucasian | 2:40 | Acute cardiac crisis | X | X | X | X |
| Sample3 | OS | AMD4 | AMD4 | NA | 94 | female | caucasian | 3:05 | COPD | X | X | X | X |
| Sample4 | OD | AMD4 | AMD4 | NA | 84 | male | caucasian | 4:23 | COPD | X | X | X | X |
| Sample5 | NA | AMD4 | AMD4 | NA | 85 | female | caucasian | 4:59 | COPD |  | X |  |  |
| Sample6 | OS | AMD3 | 4 | AMD3L | 75 | female | caucasian | 5:35 | Encephalopathy | X | X | X | X |
| Sample7 | OS | AMD3 | 4 | AMD3L | 76 | male | caucasian | 4:55 | COPD | X | X | X | X |
| Sample8 | OD | AMD3 | 4 | AMD3L | 90 | female | caucasian | 4:50 | CHF | X |  |  |  |
| Sample9 | OS | AMD3 | 4 | AMD3L | 78 | male | caucasian | 6:02 | COPD | X |  |  |  |
| Sample10 | OD | AMD3 | 4 | AMD3L | 72 | male | caucasian | 5:39 | COPD | X |  | X | X |
| Sample11 | OD | AMD3 | 4 | AMD3L | 88 | female | caucasian | 6:04 | CHF | X | X | X | X |
| Sample12 | OS | AMD3 | 4 | AMD3L | 87 | male | caucasian | 1:32 | Acute cardiac crisis | X | X | X | X |
| Sample13 | OS | AMD3 | 4 | AMD3L | 81 | male | caucasian | 5:04 | Bladder cancer | X | X | X | X |
| Sample14 | OS | AMD3 | 4 | AMD3L | 83 | male | caucasian | 3:35 | Acute cardiac crisis | X | X | X | X |
| Sample15 | OD | AMD2 | 3 | NA | 88 | male | caucasian | 3:47 | CHF | X | X | X | X |
| Sample16 | OD | AMD3 | 4 | AMD3L | 75 | female | caucasian | 4:25 | CVA |  |  | X |  |
| Sample17 | OD | AMD3 | 4 | AMD3L | 100 | female | caucasian | 5:10 | CHF | X | X | X | X |
| Sample18 | OS | AMD3 | 4 | AMD3L | 86 | male | caucasian | 3:16 | CAD | X | X | X | X |
| Sample19 | OS | AMD3 | 4 | AMD3L | 83 | male | caucasian | 5:24 | CHF | X | X | X | X |
| Sample20 | OS | AMD3 | 4 | AMD3L | 83 | male | caucasian | 3:55 | Acute cardiac crisis | X | X | X | X |
| Sample21 | OS | AMD3 | 4 | AMD3L | 90 | male | caucasian | 2:50 | Acute cardiac crisis | X | X | X | X |
| Sample22 | OD | AMD3 | 4 | AMD3L | 78 | male | caucasian | 5:59 | Acute cardiac event | X | X | X | X |
| Sample23 | OD | AMD3 | 4 | AMD3L | 80 | male | caucasian | 4:04 | Pneumonia | X | X | X | X |
| Sample24 | OS | AMD3 | 4 | AMD3L | 78 | male | caucasian | 5:27 | Acute cardiac event | X | X |  |  |
| Sample25 | OS | AMD3 | 5 | AMD3L | 86 | male | caucasian | 4:44 | MI | X | X | X | X |
| Sample26 | OD | AMD3 | 3 | AMD3L | 85 | male | caucasian | 4:37 | CHF |  | X | X | X |
| Sample27 | OD | AMD3 | 5 | AMD3L | 91 | female | caucasian | 5:56 | Acute cardiac crisis | X | X | X | X |
| Sample28 | OS | AMD3 | 5 | AMD3L | 75 | female | caucasian | 4:25 | CVA | X | X |  | X |
| Sample29 | OD | AMD3 | 5 | AMD3L | 97 | female | black | 3:52 | CHF | X | X | X | X |
| Sample30 | OD | AMD3 | 5 | AMD3L | 81 | male | caucasian | 4:40 | NA | X | X | X | X |
| Sample31 | OD | AMD3 | 5 | AMD3L | 82 | male | caucasian | 4:18 | NA | X | X | X | X |
| Sample32 | OD | AMD3 | 5 | AMD3L | 89 | female | caucasian | 5:18 | Renal failure | X | X | X | X |
| Sample33 | OS | AMD3 | 6 | AMD3M | 86 | male | caucasian | 2:32 | Acute cardiac crisis | X |  |  |  |
| Sample34 | OS | AMD3 | 6 | AMD3M | 86 | male | caucasian | 3:05 | Acute cardiac crisis | X | X | X | X |
| Sample35 | OS | AMD3 | 6 | AMD3M | 80 | female | caucasian | 2:57 | Acute cardiac crisis | X | X | X | X |
| Sample36 | OS | AMD3 | 6 | AMD3M | 86 | male | caucasian | 5:34 | CVA | X | X | X | X |
| Sample37 | OD | AMD3 | 3 | AMD3L | 82 | male | caucasian | 5:49 | Acute cardiac crisis | X | X | X | X |
| Sample38 | OD | AMD3 | 6 | AMD3M | 76 | female | caucasian | 3:44 | Pneumonia | X | X |  |  |
| Sample39 | OS | AMD3 | 7 | AMD3M | 90 | male | caucasian | 4:10 | GI bleed | X | X | X | X |
| Sample40 | OS | AMD3 | 7 | AMD3M | 79 | female | caucasian | 4:49 | Myocardial infarction | X | X | X | X |
| Sample41 | OS | AMD3 | 8 | AMD3H | 95 | female | caucasian | 4:40 | Cardiac arrest | X | X | X | X |
| Sample42 | OD | AMD3 | 8 | AMD3H | 85 | female | caucasian | 2:53 | CHF | X | X | X | X |
| Sample43 | OD | AMD3 | 6 | AMD3M | 89 | male | caucasian | 3:51 | MI | X | X | X | X |
| Sample44 | OS | AMD3 | 8 | AMD3H | 79 | male | caucasian | 4:38 | Lung cancer | X | X | X | X |
| Sample45 | OS | AMD3 | 9 | AMD3H | 85 | female | caucasian | 3:36 | Pneumonia | X | X | X | X |
| Sample46 | OD | AMD3 | 9 | AMD3H | 87 | male | caucasian | 5:39 | CHF | X | X | X | X |
| Sample47 | OS | AMD3 | 9 | AMD3H | 76 | male | caucasian | 4:21 | COPD | X | X | X | X |
| Sample48 | OS | AMD3 | 5 | AMD3L | 94 | male | caucasian | 3:22 | Acute cardiac crisis |  |  | X | X |
| Sample49 | OS | AMD3 | 9 | AMD3H | 73 | female | caucasian | 4:36 | Leukemia | X | X | X | X |
| Sample50 | OD | AMD3 | 9 | AMD3H | 85 | male | caucasian | 3:47 | Acute cardiac crisis | X |  |  |  |
| Sample51 | OD | AMD3 | 9 | AMD3H | 84 | male | black | 3:38 | Abdominal aortic aneurysm | X | X | X | X |
| Sample52 | OS | AMD3 | 9 | AMD3H | 79 | female | caucasian | 4:24 | Acute cardiac crisis | X |  |  |  |
| Sample53 | OD | AMD3 | 9 | AMD3H | 72 | male | caucasian | 2:45 | Multiple myeloma |  | X | X | X |
| Sample54 | OD | AMD3 | 9 | AMD3H | 83 | male | caucasian | 5:22 | Lymphoma | X | X | X | X |
| Sample55 | OD | AMD3 | 9 | AMD3H | 80 | female | caucasian | 5:25 | Renal failure | X | X | X | X |
| Sample56 | OD | AMD3 | 9 | AMD3H | 85 | female | caucasian | 3:27 | Necrotic bowel | X | X | X | X |
| Sample57 | OD | AMD3 | 9 | AMD3H | 85 | male | caucasian | 5:24 | Fever, severe sepsis | X | X | X | X |
| Sample58 | OD | AMD3 | 9 | AMD3H | 83 | female | caucasian | 2:39 | Multi-system failure | X | X | X | X |
| Sample59 | OD | AMD2 | 4 | NA | 79 | female | caucasian | 6:58 | Sepsis | X | X | X | X |
| Sample60 | OD | AMD3 | 9 | AMD3H | 86 | male | caucasian | 6:00 | Pneumonia | X | X | X | X |
| Sample61 | OS | AMD1 | 1 | AMD1 | 84 | male | caucasian | 2:30 | Multi-system failure | X | X | X | X |
| Sample62 | OS | AMD1 | 1 | AMD1 | 77 | female | black | 5:26 | COPD | X | X | X | X |
| Sample63 | OD | AMD1 | 1 | AMD1 | 83 | male | caucasian | 4:12 | Abdominal aortic aneurysm | X | X | X | X |
| Sample64 | OS | AMD1 | 1 | AMD1 | 83 | female | caucasian | 4:35 | Pneumonia | X | X | X | X |
| Sample65 | OS | AMD1 | 1 | AMD1 | 81 | male | caucasian | 5:54 | Acute cardiac crisis | X | X | X | X |
| Sample66 | OS | AMD1 | 1 | AMD1 | 83 | male | caucasian | 4:18 | Pneumonia | X | X | X | X |
| Sample67 | OD | AMD1 | 1 | AMD1 | 79 | female | caucasian | 3:44 | Acute cardiac crisis | X | X | X | X |
| Sample68 | OS | AMD1 | 1 | AMD1 | 82 | male | caucasian | 4:13 | CVA | X | X |  |  |
| Sample69 | OS | AMD1 | 1 | AMD1 | 78 | male | black | 4:26 | Colon cancer |  | X | X | X |
| Sample70 | OD | AMD2 | 4 | NA | 84 | male | caucasian | 4:53 | Endocarditis | X | X | X | X |
| Sample71 | OD | AMD1 | 1 | AMD1 | 83 | female | caucasian | 3:56 | MI | X | X | X | X |
| Sample72 | OD | AMD1 | 1 | AMD1 | 79 | female | caucasian | 5:10 | Pneumonia | X | X | X | X |
| Sample73 | OD | AMD1 | 1 | AMD1 | 79 | female | caucasian | 5:36 | CHF | X | X | X | X |
| Sample74 | OD | AMD1 | 1 | AMD1 | 78 | male | caucasian | 5:32 | Small bowel obstruction | X | X | X | X |
| Sample75 | OD | AMD1 | 1 | AMD1 | 79 | female | caucasian | 4:25 | Pneumonia | X | X | X | X |
| Sample76 | OD | AMD1 | 1 | AMD1 | 83 | female | caucasian | 7:00 | MI | X | X | X | X |
| Sample77 | OD | AMD1 | 1 | AMD1 | 79 | female | caucasian | 4:05 | Pneumonia | X | X | X | X |
| Sample78 | OD | AMD1 | 1 | AMD1 | 76 | female | caucasian | 4:17 | AAA |  | X | X | X |
| Sample79 | OD | AMD1 | 1 | AMD1 | 77 | male | caucasian | 4:40 | Laryngeal cancer | X | X | X | X |
| Sample80 | OD | AMD1 | 1 | AMD1 | 79 | female | caucasian | 5:19 | MI | X | X | X | X |
| Sample81 | OS | AMD3 | 4 | AMD3L | 89 | female | caucasian | 5:34 | Lung cancer | X | X | X | X |
| Sample82 | OD | AMD1 | 1 | AMD1 | 83 | male | caucasian | 5:20 | Respiratory Failure | X | X | X | X |
| Sample83 | OD | AMD1 | 1 | AMD1 | 78 | female | caucasian | 4:58 | Acute cardiac crisis | X | X | X | X |
| Sample84 | OD | AMD1 | 1 | AMD1 | 82 | female | caucasian | 4:08 | Cardiopulmonary arrest | X | X | X | X |
| Sample85 | OD | AMD1 | 1 | AMD1 | 84 | male | caucasian | 2:52 | Acute Cardiac Crisis | X | X | X | X |
| Sample86 | OD | AMD1 | 1 | AMD1 | 80 | female | caucasian | 3:35 | MI |  |  | X | X |
| Sample87 | OS | AMD1 | 1 | AMD1 | 89 | male | black | 4:43 | MI |  |  | X | X |
| Sample88 | OD | AMD1 | 1 | AMD1 | 92 | female | caucasian | 4:18 | Colitis | X | X | X | X |
| Sample89 | OS | AMD1 | 1 | AMD1 | 86 | male | caucasian | 3:36 | Pneumonia | X | X | X | X |
| Sample90 | OS | AMD2 | 2 | NA | 74 | female | caucasian | 3:53 | CHF | X | X | X | X |
| Sample91 | OS | AMD2 | 2 | NA | 84 | male | caucasian | 4:08 | Coronary artery disease | X | X | X | X |
| Sample92 | OS | AMD3 | 4 | AMD3L | 78 | female | caucasian | 4:45 | MI | X | X | X | X |
| Sample93 | OD | AMD2 | 2 | NA | 92 | male | caucasian | 3:37 | Cardiopulmonary arrest | X | X | X | X |
| Sample94 | OD | AMD2 | 2 | NA | 88 | female | caucasian | 4:03 | Sepsis | X | X | X | X |
| Sample95 | OD | AMD2 | 2 | NA | 89 | male | caucasian | 5:01 | Pneumonia | X | X | X | X |
| Sample96 | OS | AMD2 | 2 | NA | 88 | male | caucasian | 3:50 | Colon cancer | X |  | X | X |
| Sample97 | OS | AMD2 | 2 | NA | 87 | male | caucasian | 3:33 | NA | X | X | X | X |
| Sample98 | OD | AMD4 | AMD4 | NA | 74 | female | caucasian | 2:00 | Cardiac arrest | X | X | X | X |
| Sample99 | OD | AMD4 | AMD4 | NA | 88 | male | caucasian | 3:06 | Renal failure |  | X | X | X |
| Sample100 | OD | AMD4 | NA | NA | 95 | male | caucasian | 3:13 | Sepsis | X | X | X | X |
| Sample101 | OD | AMD4 | AMD4 | NA | 84 | female | caucasian | 5:09 | CVA | X | X | X | X |
| Sample102 | OD | AMD4 | AMD4 | NA | 76 | male | caucasian | 4:21 | COPD | X | X | X | X |

Abbreviations: AAA: Abdominal aortic aneurysm; CAD: Coronary Artery Disease; COPD: Chronic Obstructive Pulmonary Disorder; CHF: Congestive Heart Failure; CVA: CerebroVascular Accident; DTP: Death to Preservation time; MI: Myocardial Infarction; MC: Macular RPE/choroid; MR: Macular retina; PC: Peripheral RPE/choroid; PR: Peripheral retina. X indicates a sample of this type was used to generate the dataset.

**Supplemental Table 4.** Numbers of differentially expressed genes (DEGs) across AMD stage compared to AMD1 in bulk RNAseq.

| Tissue | DEG | AMD2 | AMD3L | AMD3M | AMD3H | AMD4 |
| --- | --- | --- | --- | --- | --- | --- |
| MC | Up | 0 | 0 | 0 | 0 | 1350 |
|  | Down | 0 | 0 | 0 | 0 | 235 |
| MR | Up | 0 | 0 | 0 | 0 | 1358 |
|  | Down | 0 | 0 | 0 | 0 | 2191 |
| PC | Up | 0 | 0 | 0 | 0 | 191 |
|  | Down | 0 | 0 | 0 | 0 | 31 |
| PR | Up | 0 | 0 | 0 | 0 | 0 |
|  | Down | 0 | 0 | 0 | 0 | 0 |

Abbreviations: MC: Macular RPE/choroid; MR: Macular retina; PC: Peripheral RPE/choroid; PR: Peripheral retina.

**Supplemental Table 5.**
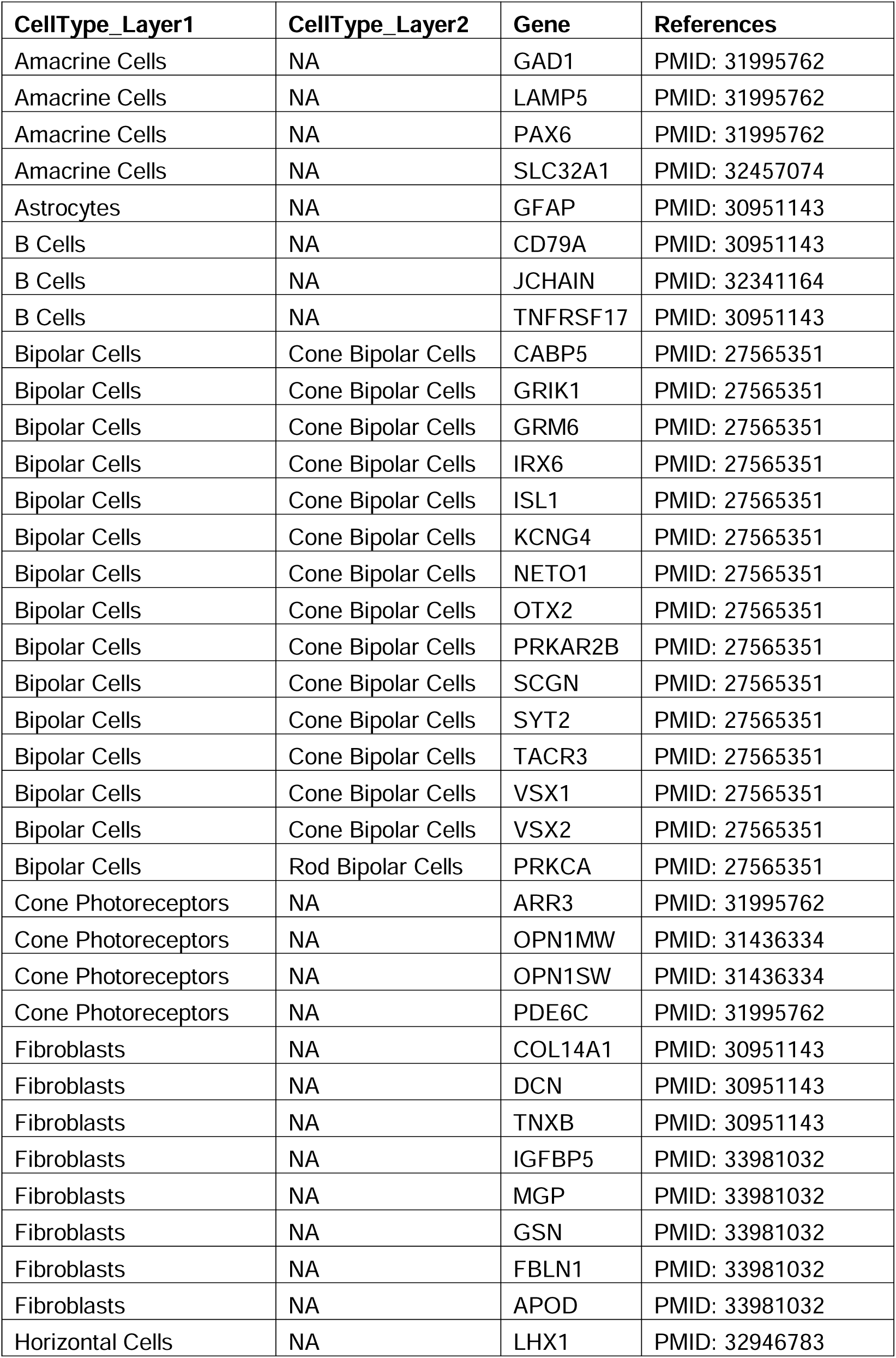
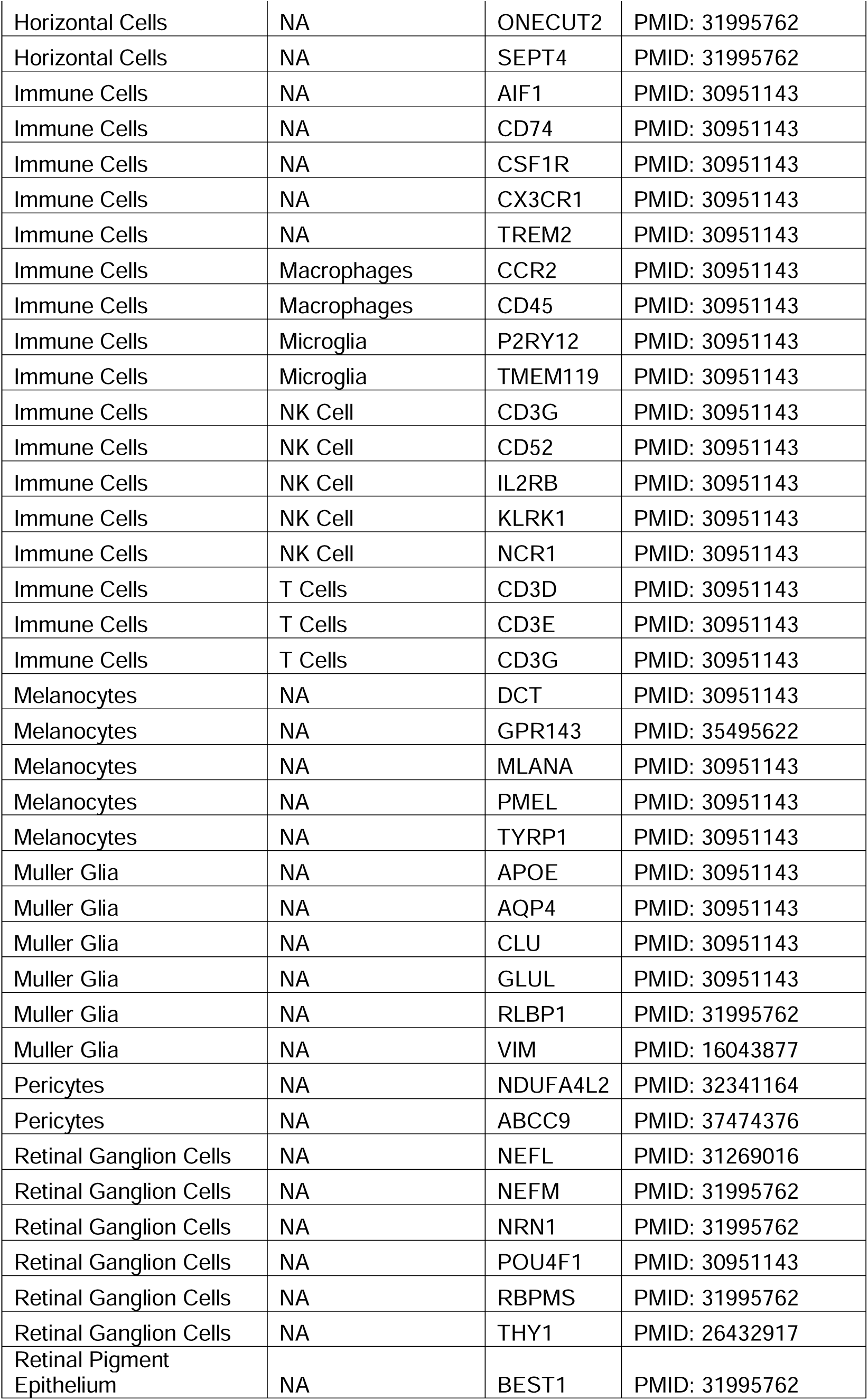
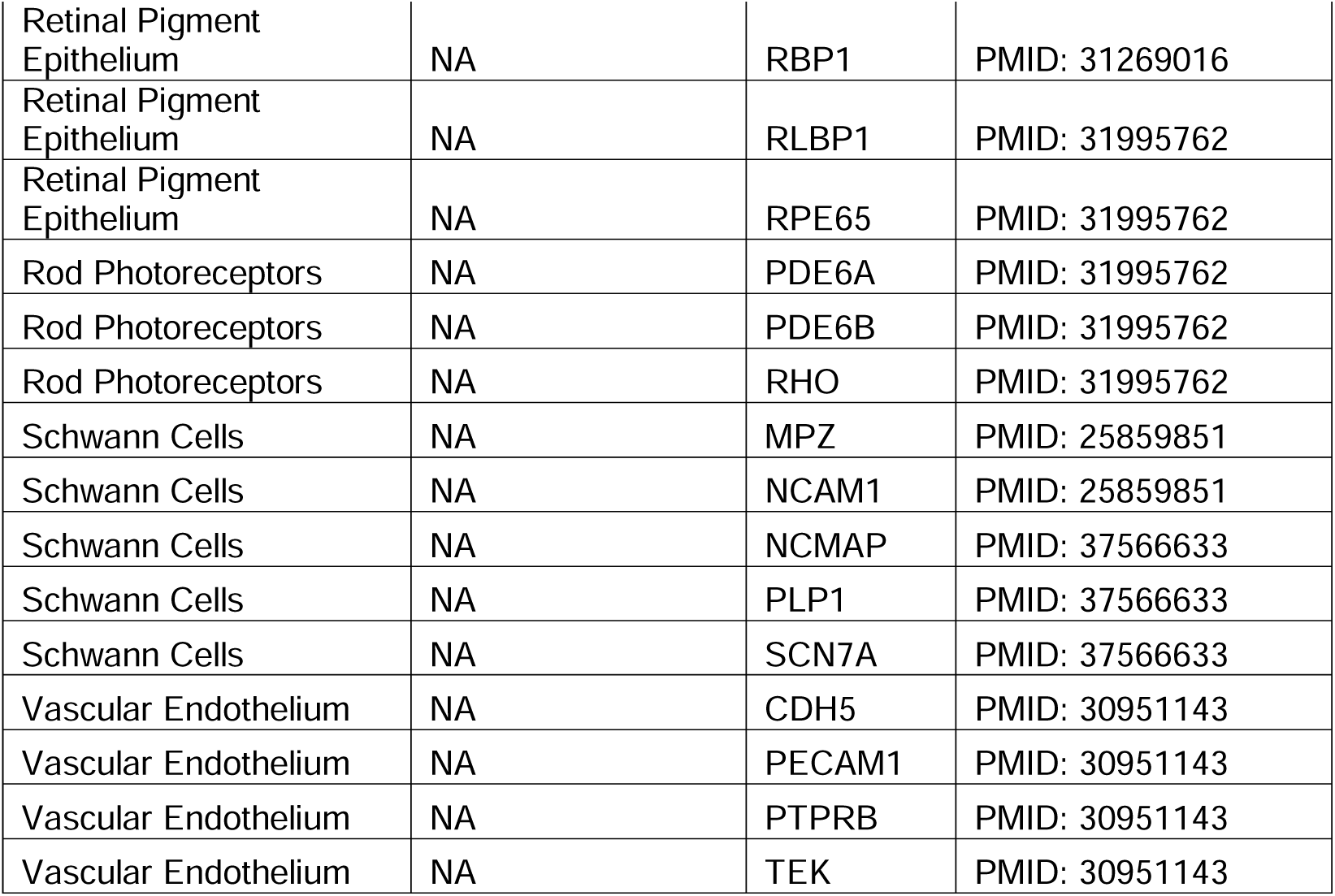
Genes used for cell type classification.

**Supplemental Table 6A.** Donor information for histology.

| Donor ID | Eye | AMD grade | Age | Sex | Race | DTP (H:MM) | Cause of Death / Recent Medical Hx |
| --- | --- | --- | --- | --- | --- | --- | --- |
| Sample H1 | OS | AMD1 | 78 | male | caucasian | 3:57 | Lung cancer |
| Sample H2 | OS | AMD1 | 67 | male | caucasian | 2:43 | Multi-system failure |
| Sample H3 | OS | AMD1 | 76 | male | caucasian | 3:46 | Liver cancer |
| Sample H4 | OS | AMD1 | 85 | male | caucasian | 4:46 | Cardiopulmonary arrest |
| Sample H5 | OS | AMD1 | 80 | male | black | 3:53 | Sepsis |
| Sample H6 | OS | AMD1 | 56 | male | caucasian | 3:41 | Hypoxia |
| Sample H7 | OS | AMD1 | 88 | male | caucasian | 2:41 | GI bleed |
| Sample H8 | OS | AMD1 | 73 | male | caucasian | 5:37 | Prostate cancer |
| Sample H9 | OS | AMD1 | 73 | male | caucasian | 6:08 | Leukemia |
| Sample H10 | OS | AMD1 | 86 | male | caucasian | 5:59 | Pneumonia |
| Sample H11 | OS | AMD1 | 81 | male | caucasian | 6:27 | CVA |
| Sample H12 | OS | AMD1 | 93 | female | caucasian | 5:59 | CVA |
| Sample H13 | OS | AMD1 | 75 | male | caucasian | 3:29 | Diabetes mellitus |
| Sample H14 | OS | AMD1 | 79 | female | caucasian | 5:08 | COPD/CKD |
| Sample H15 | OS | AMD2 | 72 | female | caucasian | 4:06 | CHF |
| Sample H16 | OS | AMD2 | 64 | female | caucasian | 3:58 | Leukemia |
| Sample H17 | OS | AMD2 | 73 | male | caucasian | 5:38 | Aortic stenosis |
| Sample H18 | OS | AMD2 | 84 | male | caucasian | 3:17 | Sepsis |
| Sample H19 | OS | AMD2 | 76 | female | caucasian | 5:17 | Acute cardiac event |
| Sample H20 | OS | AMD2 | 97 | female | caucasian | 4:19 | Acute cardiac event |
| Sample H27 | OS | AMD3H | 81 | female | caucasian | 4:43 | CHF |
| Sample H28 | OS | AMD3H | 97 | male | caucasian | 5:30 | Acute cardiac event |
| Sample H29 | OS | AMD3H | 90 | female | caucasian | 3:02 | GI bleed |
| Sample H30 | OS | AMD3H | 73 | female | caucasian | 5:17 | Multi-system failure |
| Sample H31 | OS | AMD3H | 78 | male | caucasian | 5:59 | Acute cardiac event |
| Sample H32 | OS | AMD3H | 80 | male | caucasian | 3:35 | Acute cardiac event |
| Sample H34 | OS | AMD3H | 80 | female | caucasian | 4:11 | Myocardial infarction |
| Sample H35 | OS | AMD3H | 94 | female | caucasian | 4:06 | CHF |
| Sample H36 | OS | AMD3L | 79 | female | caucasian | 2:24 | Lung cancer |
| Sample H37 | OS | AMD3L | 76 | male | caucasian | 4:32 | Lung cancer |
| Sample H38 | OS | AMD3L | 76 | female | caucasian | 6:27 | Sepsis |
| Sample H39 | OS | AMD3L | 72 | male | caucasian | 4:57 | Respiratory failure |
| Sample H40 | OS | AMD3L | 90 | female | caucasian | 4:18 | Failure to thrive |
| Sample H42 | OS | AMD3L | 72 | male | caucasian | 2:55 | Acute cardiac event |
| Sample H43 | OS | AMD3L | 75 | male | caucasian | 5:06 | Pancreatitis |
| Sample H44 | OS | AMD3L | 89 | female | caucasian | 5:18 | Renal failure |
| Sample H45 | OS | AMD3M | 89 | male | caucasian | 3:34 | CHF |
| Sample H46 | OS | AMD3M | 97 | female | caucasian | 4:33 | Pleural effusion |
| Sample H47 | OS | AMD3M | 88 | male | caucasian | 3:16 | CHF |
| Sample H48 | OS | AMD3M | 90 | male | caucasian | 4:07 | CKD |
| Sample H49 | OS | AMD4 | 90 | male | caucasian | 3:55 | CHF |
| Sample H50 | OS | AMD4 | 93 | female | caucasian | 3:15 | Multi-system failure |
| Sample H51 | OS | AMD4 | 80 | male | caucasian | 2:24 | Sepsis |
| Sample H52 | OS | AMD4 | 81 | male | caucasian | 4:09 | GI bleed |
| Sample H53 | OS | AMD4 | 79 | male | caucasian | 3:50 | Respiratory failure |
| Sample H54 | OS | AMD4 | 94 | female | caucasian | 3:48 | CHF |

Abbreviations: CKD: Chronic Kindey Disease; COPD: Chronic Obstructive Pulmonary
Disorder; CHF: Congestive Heart Failure; CVA: CerebroVascular Accident; DTP: Death to
Preservation time.

**Supplemental Table S6B.** Number of donors with histology stains by AMD grade.

|  | H&E | IBA1 | GFAP | C3 | TREM2 | OLR1 | NGFR | SPP1 | GPX3 |
| --- | --- | --- | --- | --- | --- | --- | --- | --- | --- |
| AMD1 | 14 | 6 | 2 | 4 | 4 | 4 | 5 | 6 | 5 |
| AMD2 | 6 | 4 | 2 | 0 | 0 | 0 | 0 | 0 | 0 |
| AMD3L | 8 | 1 | 0 | 6 | 4 | 5 | 5 | 6 | 6 |
| AMD3M | 4 | 1 | 3 | 0 | 0 | 0 | 0 | 0 | 0 |
| AMD3H | 8 | 2 | 1 | 6 | 6 | 5 | 4 | 6 | 6 |
| AMD4 | 6 | 5 | 1 | 5 | 5 | 4 | 4 | 5 | 5 |
| **Total** | 46 | 19 | 9 | 21 | 19 | 18 | 18 | 23 | 22 |

**Supplemental Table 7:** Donor demographics for samples used in TREM2 ELISA.

| **MGS4** | **Sex** | **Age** | **Race** | **DTP**<br>**(H:MM)** | **Cause of Death /**<br>**Recent Medical Hx** | **Serum** | **AqH** | **VH** | **MC** | **PC** | **MR** | **PR** |
| --- | --- | --- | --- | --- | --- | --- | --- | --- | --- | --- | --- | --- |
| AMD1 | female | 90 | Caucasian | 5:25 | Cardiac | X | X | X | X | X | X | X |
| AMD1 | male | 86 | Caucasian | 4:47 | CHF | X | X | X | X | X | X | X |
| AMD1 | female | 93 | Caucasian | 4:10 | GI Bleed | X | X | X | X | X | X | X |
| AMD1 | male | 83 | Caucasian | 6:03 | CVA | X | X | X | X | X | X | X |
| AMD1 | male | 73 | Caucasian | 6:10 | Cardiac | X | X | X | X | X | X | X |
| AMD1 | male | 85 | Caucasian | 3:56 | CHF | X | X | X | X | X | X | X |
| AMD1 | male | 89 | Caucasian | 3:05 | Cardiac | X | X | X | X | X | X | X |
| AMD1 | female | 75 | Caucasian | 5:52 | CFH | X | X | X | X | X | X | X |
| AMD4 | female | 75 | Caucasian | 2:36 | Perforated<br>diverticulitis | NA | X | X | X | X | X | X |
| AMD4 | male | 92 | Caucasian | 6:15 | Cardiac | X | X | X | X | X | X | X |
| AMD4 | male | 84 | Caucasian | 3:29 | Respiratory | X | X | X | X | X | X | X |
| AMD4 | female | 91 | Caucasian | 3:58 | Cardiac | NA | X | X | X | X | X | X |
| AMD4 | female | 94 | Caucasian | 5:25 | NA | X | X | X | X | X | X | X |
| AMD4 | male | 74 | Caucasian | 4:26 | COPD | X | X | X | X | X | X | X |
| AMD4 | male | 86 | Caucasian | 5:26 | Cardiac | X | X | X | X | X | X | X |
| AMD4 | male | 81 | Caucasian | 4:55 | ESRD | X | X | X | X | X | X | X |

Abbreviations: AqH: Aqueous Humor; DTP: Death to Preservation Time; MC: Macular
RPE/choroid; MR: Macular retina; PC: Peripheral RPE/choroid; PR: Peripheral retina; VH:
Vitreous Humor. X indicates a sample of this type was used to generate the dataset.

